# The Hippo Signaling Pathway Regulates Corneal Endothelial Regeneration

**DOI:** 10.64898/2025.12.30.697046

**Authors:** Jingbin Zhuang, Yuli Guo, Houjian Zhang, Meiqin Zhong, Xu Hu, Lingyu Zhang, Jingwen Yu, Yuqian Wang, Shundong Cai, Huping Wu, Chengchao Chu, Hui He, Rongrong Zong, Weifu Huang, Andrew J. Quantock, Zuguo Liu, Wei Li

## Abstract

Corneal endothelial cells (CECs) have limited regeneration capacity in primates while display a potent restorative capability in some species such as rodents after wounding. The mechanism which determines the regenerative capability of CECs remains poorly understood. Here, we identify that the Hippo signaling pathway is inhibited and the downstream effector YAP is activated during CEC wound healing in rabbit and mouse. Knockdown of *Yap1* and pharmacological inhibition of YAP suppresses CECs wound healing in rodents in both *in vitro* and *in vivo* models. XMU-MP-1, a specific small molecular inhibitor for Hippo signaling pathway, can promote the corneal endothelial regeneration in rodents and corneal endothelial proliferation in cultured primate CECs, as well as *in vivo* non-human primate CEC wounding model. These findings suggest that Hippo pathway serves as a conserved signal to regulate corneal endothelial regeneration among various species, providing a novel target for non-invasive treatment of corneal endothelium decompensation.

## 1. Introduction

The inner surface of cornea consists a single layer of hexagonal corneal endothelial cells (CECs) which maintain hydration and transparency of cornea through barrier function and pump function of CECs.^[1, 2]^ The normal density of CEC in adult humans is 2500 cells/mm^2^ in average.^[3]^ When CEC density falls below a critical threshold around 500 cells/mm², the corneal endothelial function will be decompensated, with the result of corneal edema, opacity and progressive vision loss ^[4, 5]^, constituting one of the leading causes of corneal blindness.^[6]^

Currently, surgical treatment is the major therapeutic approach for corneal endothelial decompensation, including Descemet stripping endothelial keratoplasty and Descemet’s membrane endothelial keratoplasty.^[7, 8]^ However, it is limited by global shortage of corneal donor tissue. Several researches tried to transplant cultured human CECs ^[9, 10]^ and human pluripotent stem cell derived CECs ^[11]^ to replenish the lost CECs. A series of studies on animal models ^[12–15]^ and clinical trials ^[16–18]^ have shown that Rho-kinase inhibitors (ROCKi) could promote CEC wound healing. However, there are currently no pharmaceutical interventions clinically approved for corneal endothelial decompensation.

The regenerative capacity of CEC varies among species. Although CECs in primates exhibit limited *in vivo* proliferation capacity, CECs in rodents such as rabbit and mouse demonstrate a robust ability to regenerate,^[19, 20]^ while underneath mechanism is poorly understood. Over the past few years, tissues from other species possessing regenerative capability have been extensively utilized to study the molecular mechanism, contributing to identification of novel pharmaceutical targets to treat human diseases.^[21–23]^ Therefore, investigating the mechanism of corneal endothelial wound healing in rodents may provide insights for exploring strategies for human corneal endothelial regeneration.

The Hippo pathway has been reported to play a crucial role in tissue wound healing and organ development and regeneration.^[24–27]^ It functions via a phosphorylation cascade involving MST1/2, LATS1/2, and MOB1A/B proteins, which subsequently phosphorylates the downstream transcriptional cofactors YAP/TAZ for their cytoplasmic retention. When the Hippo pathway is inactivated, unphosphorylated YAP/TAZ translocates into the nucleus and initiates transcription of pro-proliferative and pro-survival genes to promote cell regeneration.^[28, 29]^ However, whether this evolutionarily conserved pathway contributes to the regenerative capacity of CECs remains unknown. In this study, we investigated the role of Hippo signaling pathway on corneal endothelial wound healing and determined whether this signaling pathway can be a regulator of corneal endothelial regeneration in multiple species.

## 2. Results

### 2.1 Hippo-YAP pathway changes during corneal endothelial wound healing

To determine whether the Hippo-YAP signaling axis is engaged in CEC wound healing, we investigated the status of Hippo pathway during CEC wound healing across species using a rabbit CEC cryoinjury model (**Figure 1**A), a mouse ultraviolet (UV)-induced CEC damage model (Figure 1E), and an *in vitro* primary monkey CEC scraping model (Figure S1A, Supporting Information). YAP phosphorylation was significantly decreased on day 7 and returned to baseline level on day 14 after CEC cryoinjury in rabbit (Figure 1B,C). Phosphorylated MST1 and LATS1, two key kinases in the Hippo pathway, showed similar pattern (Figure 1B,C). There was obvious YAP nuclear translocation on day 7 compared to day 0 and day 14 (Figure 1D), indicating the activation of YAP. Similar finding was also detected during wound healing of UV-induced mouse CEC damage (Figure 1F) and the primary monkey CEC scraping model (Figure S1B,C, Supporting Information). Collectively, these results indicated that the Hippo pathway was downregulated and YAP was activated during CEC wound healing in different species.

**Figure 1.**
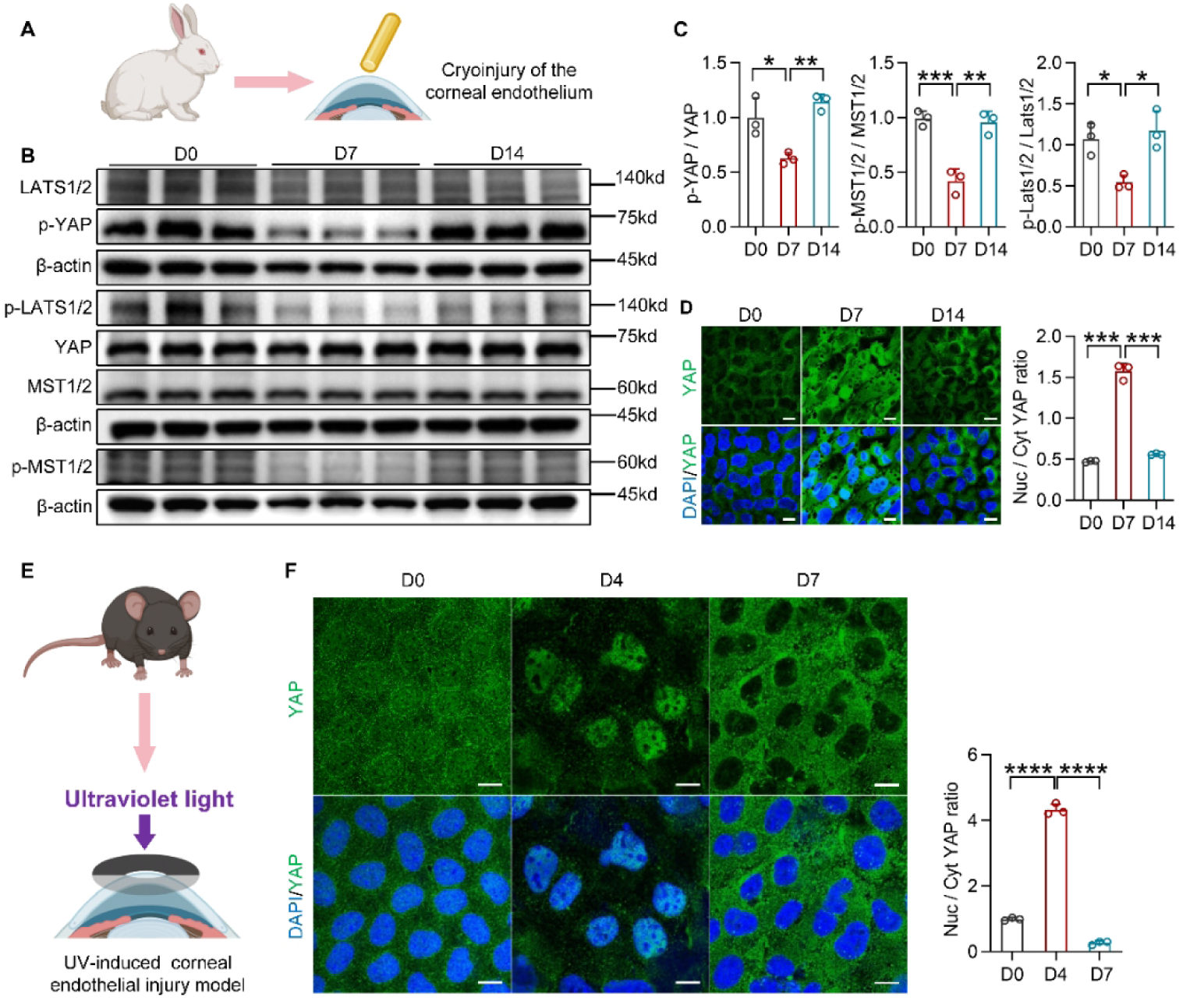
Downregulation of Hippo pathway and YAP activation during CEC wound healing. (A) The diagram shows the rabbit corneal endothelial injury model. (B, C) Western blot analysis of p-YAP, YAP, p-MST1/2, MST1/2, p-LATS1/2, LATS1/2 and β-actin, the protein levels were quantified by densitometry (n=3). (D) Representative immunofluorescence images of YAP in rabbits and the ratio of nuclear YAP to cytoplasm YAP, five cells were randomly selected for measurement and averaged in each sample (n = 3). (E) The diagram shows the mouse corneal endothelial injury model. (F) Representative immunofluorescence images of YAP in mice and the ratio of nuclear YAP to cytoplasm YAP, five cells were randomly selected for measurement and averaged in each sample (n = 3). *P < 0.05, **P < 0.01, ***P < 0.001, ****P < 0.0001, error bars, mean ± SEM. Scale bars represent 10 µm in (D) and (F). Nuc, nucleus; Cyt, cytoplasm.

### 2.2 Hippo pathway is a major regulator of CEC regeneration

To further elucidate the role of Hippo pathway in CEC regeneration, we applied verteporfin, a specific inhibitor of YAP which disrupts the YAP-TEAD interaction,^[30]^ to block YAP function. EdU labeling and Ki67 staining showed reduced proliferation of primary rabbit CECs under verteporfin treatment (**Figure 2**A). CCK8 assay showed that verteporfin inhibited viability of primary rabbit CECs in a dose-dependent manner (Figure 2B). Human CEC line B4G12 demonstrated similar changes after YAP inhibition (Figure S2, Supporting Information). Verteporfin eye drops were subsequently administrated to treat rabbits following corneal endothelial cryoinjury, and the cornea showed reduced transparency on day 7 and cloudy central cornea on day 13 (Figure 2C,F). Anterior segment optical coherence tomography (OCT) examination showed delayed recovery of corneal edema in verteporfin treated rabbit (Figure 2D,F). Alizarin red staining of rabbit corneas on day 13 confirmed delayed CEC wound healing after verteporfin application, characterized by absence of cellular structure in the central corneal endothelium (Figure 2E). Similar results were observed in mouse with UV radiation induced corneal endothelial damage (Figure 2G-J). Putting together, pharmacological YAP inhibition suppressed the regeneration of CECs in both *in vitro* and *in vivo* models across species.

**Figure 2.**
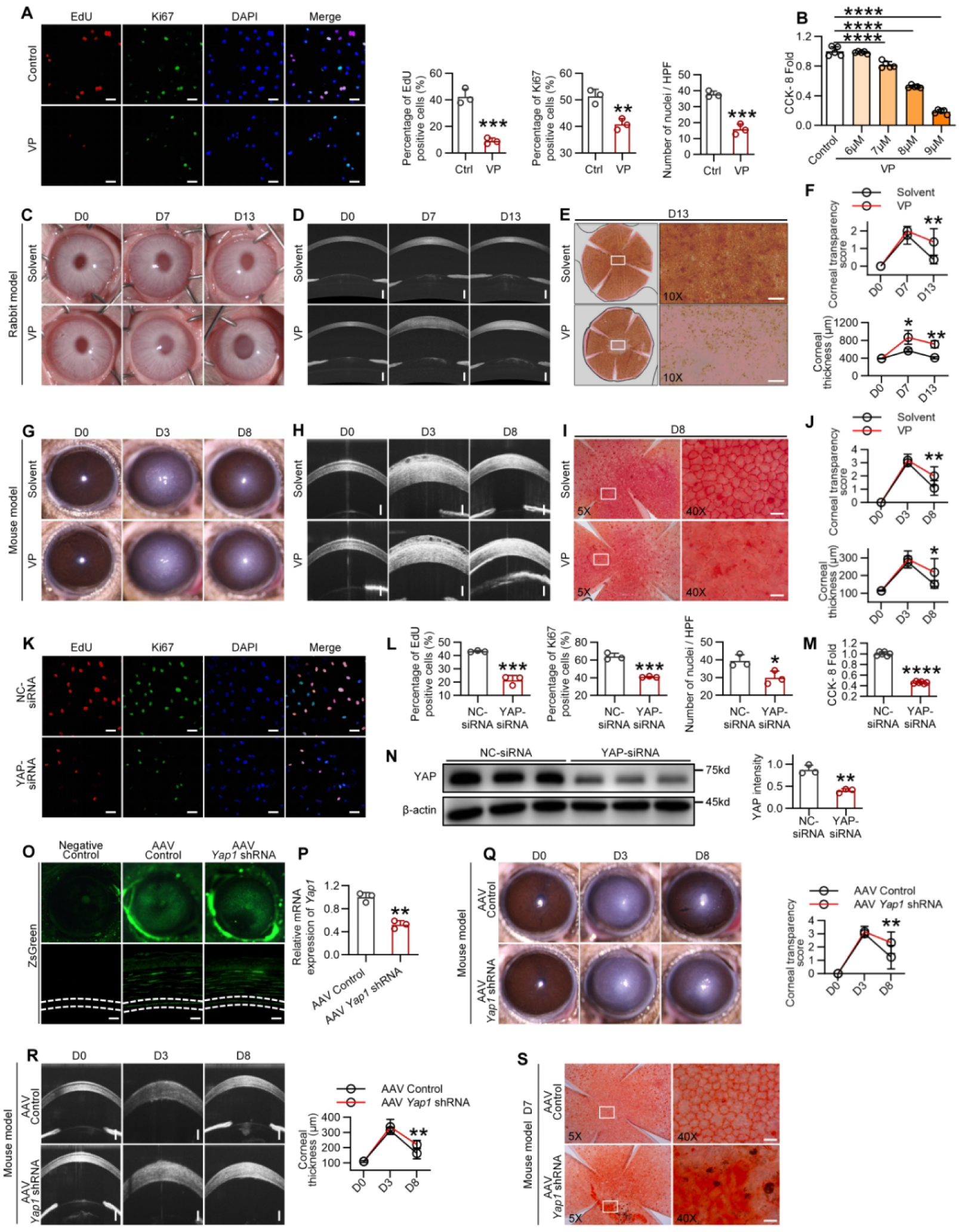
Pharmacological YAP inhibition and knockdown of Yap1 suppressed CEC regeneration. (A) Representative staining images of primary rabbit CECs and percentage of EdU and Ki67 positive cells and counting of DAPI (n = 3). (B) CCK-8 assay of primary rabbit CECs. (C to E) Representative slit-lamp images (C), anterior segment OCT images (D) and corneal alizarin red staining images (E) of rabbits. (F) Corneal transparency score and central corneal thickness of rabbits (n = 4). (G to J) Representative images (G to I) and corneal transparency score and central corneal thickness of mouse models (n = 5) (J). (K and L) Representative images (K) of EdU and Ki67 and percentage of EdU, Ki67 positive cells and counting of DAPI (n = 3) (L). (M) CCK-8 assay of primary rabbit CECs. (Fold of control, n = 6). (N) Western blot analysis of YAP in primary rabbit CECs (n = 3). (O) ZsGreen expression in vivo and in corneal sections. (P) Relative mRNA expression of Yap1 in mice corneal endothelium (n = 3). (Q) Representative slit-lamp images and corneal transparency score of mice (n = 6). (R) Representative anterior segment OCT images and central corneal thickness of mouse models (n = 6). (S) Representative corneal alizarin red staining images of mice. *P < 0.05, **P < 0.01, ***P < 0.001, ****P < 0.0001, error bars, mean ± SEM. Scale bars represent 500 µm in (D), 200 µm in (E), 100 µm in (H and R), 50 µm in (A and K), 25 µm in (O), and 20 µm in (I and S). VP, verteporfin.

To further confirm the function of YAP in corneal endothelium, primary rabbit CECs were transfected with a siRNA specific to *Yap1*. EdU and Ki67 positive cells were significantly reduced after knockdown of *Yap1* (Figure 2K,L,N). Cell viability was also reduced after knockdown of *Yap1* as measured by CCK8 assay (Figure 2M). Similar results were observed in B4G12 cell line (Figure S3, Supporting Information). Next, we determined the effect of *Yap1* knockdown on wound healing in a mouse model. The adeno-associated virus (AAV) vector carrying *Yap1*-specific short hairpin RNA (AAV-*Yap1*-shRNA) was injected into mouse anterior chamber and verified by ZsGreen expression in the corneal endothelium 4 weeks after injection (Figure 2O). *Yap1* mRNA expression was suppressed in the corneal endothelium of AAV-*Yap1*-shRNA mice (Figure 2P), and the mice showed a delayed recovery from corneal opaque and stromal edema after UV radiation damage (Figure 2Q,R,T). Taken together, pharmacological YAP inhibition and *Yap1* knockdown significantly suppressed corneal endothelial regeneration, supporting the function of Hippo signaling pathway in corneal endothelial wound healing.

### 2.3 Pharmaceutical inhibition of Hippo pathway with XMU-MP-1 promotes CEC regeneration

XMU-MP-1, a specific MST1/2 inhibitor,^[31]^ can block Hippo pathway and restrain the phosphorylation of YAP. We applied XMU-MP-1 in primary rabbit CEC culture and found it can reduce YAP phosphorylation and enhance YAP nuclear translocation in rabbit CECs (Figure S4A,C,E,F, Supporting Information). Additionally, XMU-MP-1 increased EdU and Ki67 positive cells in both rabbit CECs (**Figure 3**A,B) and human B4G12 cell line (Figure S5A-D), and enhanced the cell viability (Figure 3C, Figure S5E, Supporting Information). Cell cycle analysis revealed a significant decrease of cells in G1-phase and increase of cells in G2/M-phase after XMU-MP-1 application (Figure S6A,B, Supporting Information). XMU-MP-1 treatment also upregulated the expression of cell cycle-related genes including E2F1, CCND1, CCND3, CCNA2 and CCNE1 (Figure S6C,D, Supporting Information).

**Figure 3.**
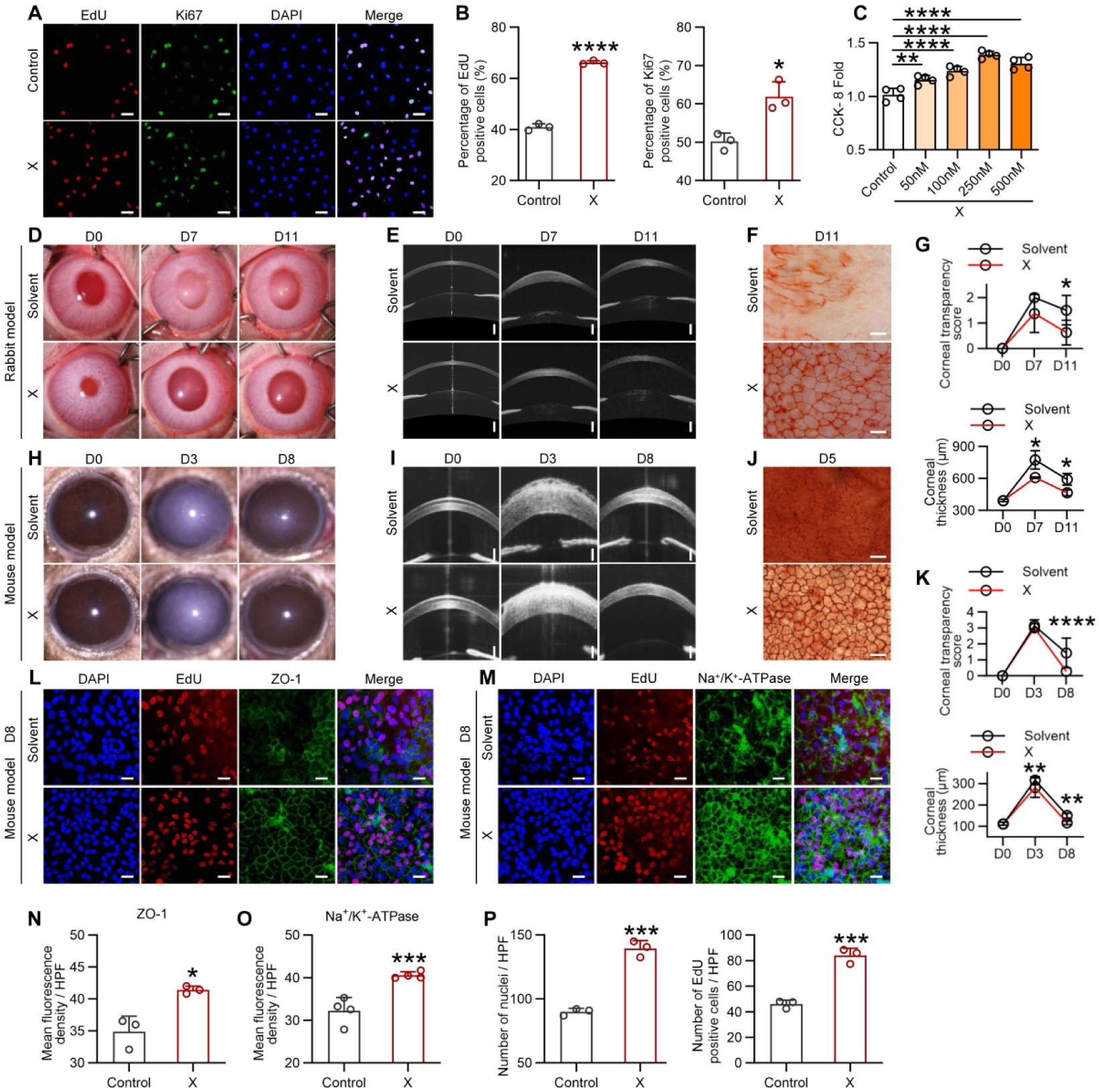
XMU-MP-1 promotes CEC regeneration in rabbit and mouse models. (A and B) Representative staining images of primary rabbit CECs and percentage of EdU and Ki67 positive cells (n = 3). (C) CCK-8 assay of primary rabbit CECs (fold of control, n=4). (D to G) Representative images (D to F) and corneal transparency score and central corneal thickness of rabbit models (n = 3) (G). (H to K) Representative images (H to J) and corneal transparency score and central corneal thickness of mouse models (n = 6) (K). (L and M) EdU staining and immunofluorescence of ZO-1 (L), Na^+^-K^+^ ATPase (M) of corneal endothelial whole mount. (N and O) The intensities of ZO-1 (N) and Na^+^-K^+^ ATPase (O) (n = 3). (P) Quantification of EdU positive cells and nuclei counting in (L and M) (n = 3). *P < 0.05, **P < 0.01, ***P < 0.001, ****P < 0.0001, error bars, mean ± SEM. Scale bars represent 500 µm in (E), 100 µm in (I), 50 µm in (A), 25 µm in (L and M) and 20 µm in (F and J). X, XMU-MP-1; ip, intraperitoneal injection.

Furthermore, XMU-MP-1 was topically applied in corneal endothelial wound models and found it expedited corneal endothelial wound healing and corneal edema recovery in both rabbits (Figure 3D-G) and mice (Figure 3H-K). Immunostaining revealed that XMU-MP-1-treated mouse corneas showed more contiguous ZO-1 (Figure 3L,N) and increased Na^+^/K^+^-ATPase (Figure 3M,O) expression. Additionally, more EdU-positive cells were found in the corneal endothelium of XMU-MP-1 treated mice (Figure 3P). These results indicated that XMU-MP-1 could promote CEC regeneration and functional restoration.

To verify whether XMU-MP-1 can penetrate the cornea, liquid chromatography-tandem mass spectrometry analysis was utilized to assay the concentration of XMU-MP-1 in the rabbit corneal endothelium 1 hour after topical application, and 2974.44 ± 17.43 ng/g of XMU-MP-1 was detected (Figure S7A, Supporting Information). The nuclear translocation of YAP in mouse CECs was also observed following administration of XMU-MP-1 (Figure S7B, Supporting Information).

We further evaluated the ocular manifestation after 4 weeks topical administration of XMU-MP-1 in normal mice. No corneal limbal neovascularization, corneal epithelial defect or corneal thickness changes were observed after XMU-MP-1 application (Figure S8A,B, Supporting Information). There was also no evidence of abnormal tissue proliferation in the conjunctiva, Meibomian glands, or lacrimal glands (Figure S8C, Supporting Information). Additionally, there was no significant difference in tear production, intraocular pressure, axial length, or refraction (Figure S8D-G, Supporting Information) compared with that of the control group. There was also no abnormal retinal thickening or neovascularization detected after XMU-MP-1 application (Figure S8H,I, Supporting Information).

### 2.4 XMU-MP-1 promotes CEC regeneration in primates

To investigate the effect of XMU-MP-1 on primate CECs, we firstly treated the primary cultured human CECs (**Figure 4**A,B) and monkey CECs (Figure 4C,D) with XMU-MP-1. The results showed that EdU-positive and Ki67-positive cells in both types of CECs increased with XMU-MP-1 administration. Nuclei counting confirmed higher cell density in XMU-MP-1 treated groups despite equal seeding densities (Figure 4A-D). Next, the XMU-MP-1 treatment was applied to an *ex vivo* monkey corneal endothelial scraping model. Trypan blue staining demonstrated that XMU-MP-1 promoted monkey CEC wound healing, with the defect area completely healed by day 4 post scraping (Figure 4E,F).

**Figure 4.**
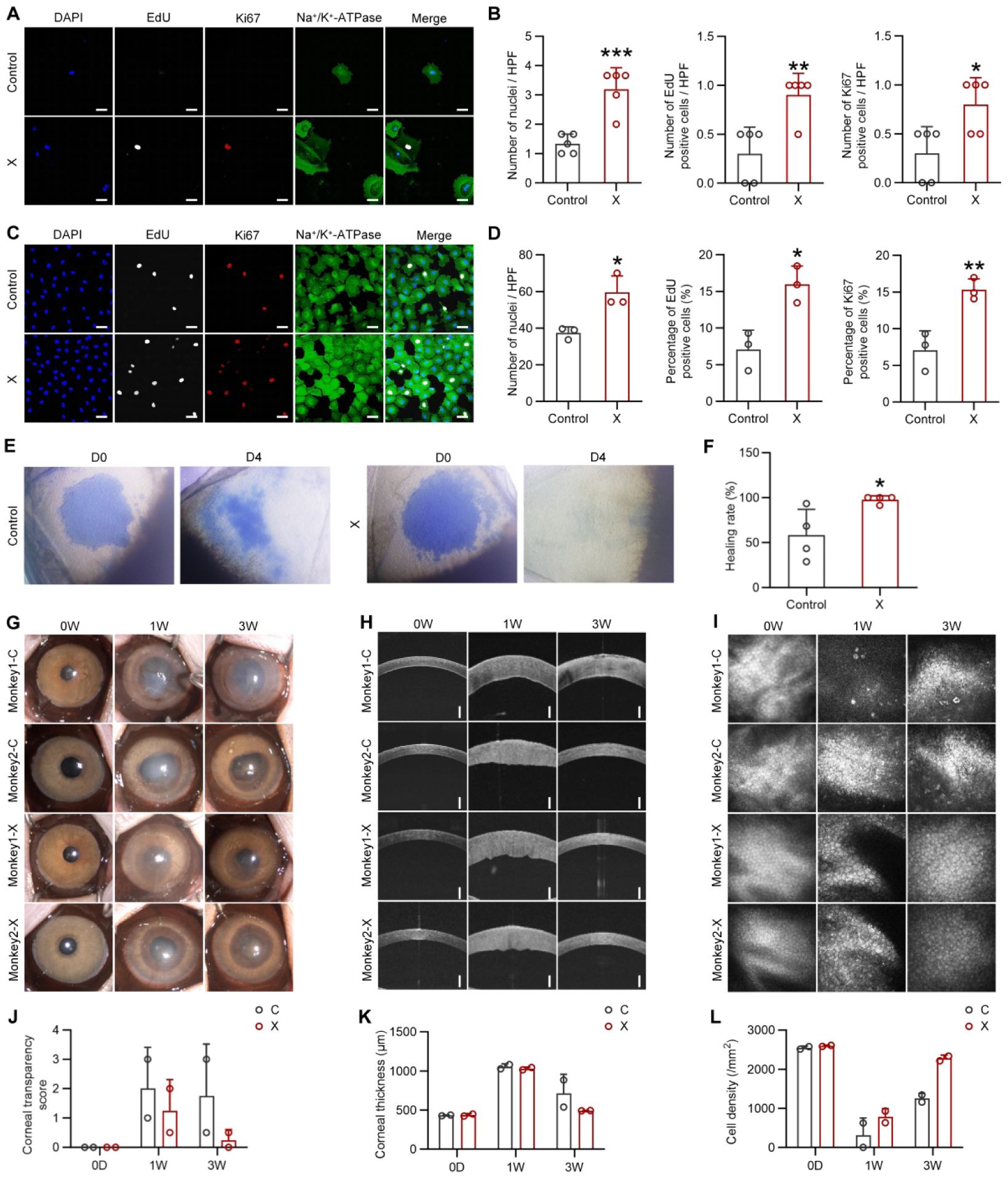
XMU-MP-1 promotes CEC regeneration in non-human primate. (A) EdU and DAPI staining and immunofluorescence of Ki67, Na^+^-K^+^ATPase in primary human CECs. (B) Nuclei counting and percentage of EdU and Ki67 positive cells in (A) (n = 5). (C) EdU and DAPI staining and immunofluorescence of Ki67 and Na^+^-K^+^ ATPase in primary monkey CECs. (D) Nuclei counting and percentage of EdU and Ki67 positive cells in (C) (n = 3). (E and F) Trypan blue staining of monkey corneal endothelium explants and the healing rate quantified by stained area (n=4). (G to I) Slit-lamp images (G), anterior segment OCT images (H) and corneal endothelial in vivo confocal microscope graphs (I) of monkeys. (J to L) Corneal transparency score (J), central corneal thickness (K) and central CECs density (L) of monkeys (n=2). *P < 0.05, **P < 0.01, ***P < 0.001, ****P < 0.0001, error bars, mean ± SEM. Scale bars represent 500 µm in (H) and 25 µm in (A and C). C, control; X, XMU-MP-1.

Next, XMU-MP-1 solution was topically applied in cynomolgus monkey after cryoinjury of the central corneas. The slit lamp microscopy (Figure 4G,J) and anterior segment OCT examination (Figure 4H,K) showed that the corneal transparency and thickness restored to normal in XMU-MP-1 treated eyes at week 3, while one control cornea remained obvious opacity and edema. CECs were observed under in vivo confocal microscopy and regular cell morphology was observed 3 weeks after topical application of XMU-MP-1 (Figure 4I). CECs density in the central cornea was restored to pre-injury level with XMU-MP-1 treatment, while it was about half of pre-injury level in control eyes (Figure 4L). These findings suggest that XMU-MP-1 effectively promotes CEC regeneration in non-human primates.

We further conducted a 12-month follow-up in one monkey after 3-weeks XMU-MP-1 treatment. Slit-lamp examination revealed clear cornea in XMU-MP-1 treated eye, while the solvent-treated control eye exhibited stromal opacification (**Figure 5**A). Corresponding anterior segment OCT confirmed lower corneal thickness in XMU-MP-1-treated eye, compared with that of control eye (Figure 5B). In vivo confocal microscopy demonstrated visible, regularly arranged CECs in XMU-MP-1-treated eye, while CECs were undetectable in control eye (Figure 5C). Alizarin red staining further revealed a confluent endothelial monolayer with uniform hexagonal morphology in XMU-MP-1-treated cornea, compared to sparse, irregular cells in control eye (Figure 5D). Additionally, CECs showed integral expression and organized localization of Na⁺/K⁺-ATPase and ZO-1 in treated cornea, while control cornea displayed disorganized expression of both markers (Figure 5E). Collectively, these results demonstrate that a short-term XMU-MP-1 treatment promoted corneal endothelial regeneration after wounding, and the structure and function of corneal endothelium could be maintained over an extended period without further therapeutic intervention.

**Figure 5.**
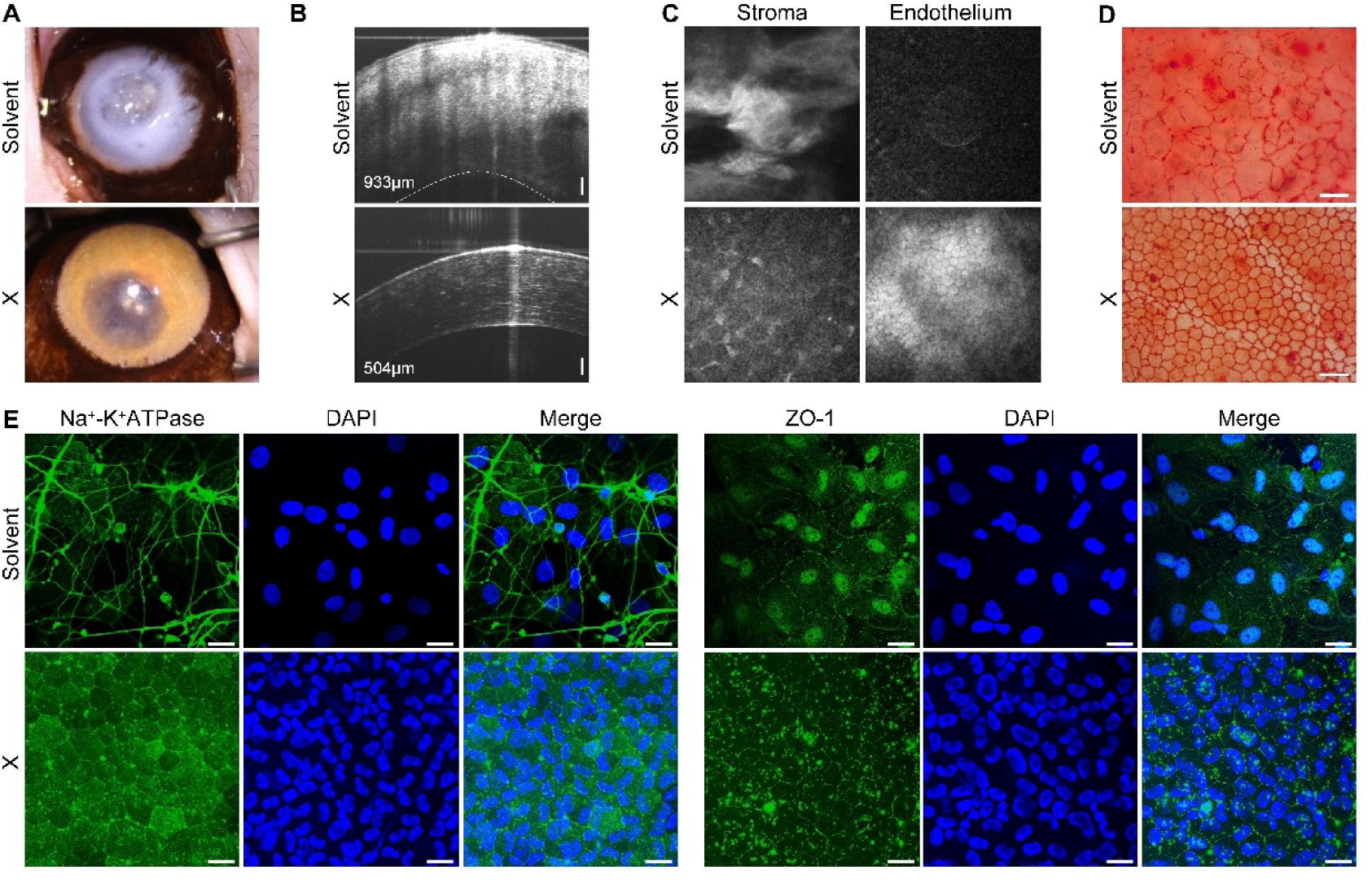
Long-term follow-up on monkey corneal endothelium transiently treated with XMU-MP-1 post-injury. (A to C) Representative slit-lamp images (A), anterior segment OCT images (B) and in vivo confocal microscope graphs (C) of XMU-MP-1-treated eye and the solvent control eye at 12 months after treatment cessation. (D) Alizarin red staining of monkey corneal endothelium. (E) Immunofluorescence staining of Na^+^/K^+^-ATPase and ZO-1 in monkey corneal endothelium. Scale bars represent 100 µm in (B), 40 µm in (D), and 25 µm in (E). X, XMU-MP-1.

## 3. Discussion

In this study, we identified the activation of Hippo-YAP signaling axis during CEC wound healing and confirmed its regulatory role in CEC regeneration. Hippo pathway inhibitor exhibited efficacy in promoting CEC regeneration across mouse, rabbit and monkey corneal endothelial wounding models (**Figure 6**), indicating the potential of its clinical application in treating corneal endothelial decompensation.

**Figure 6.**
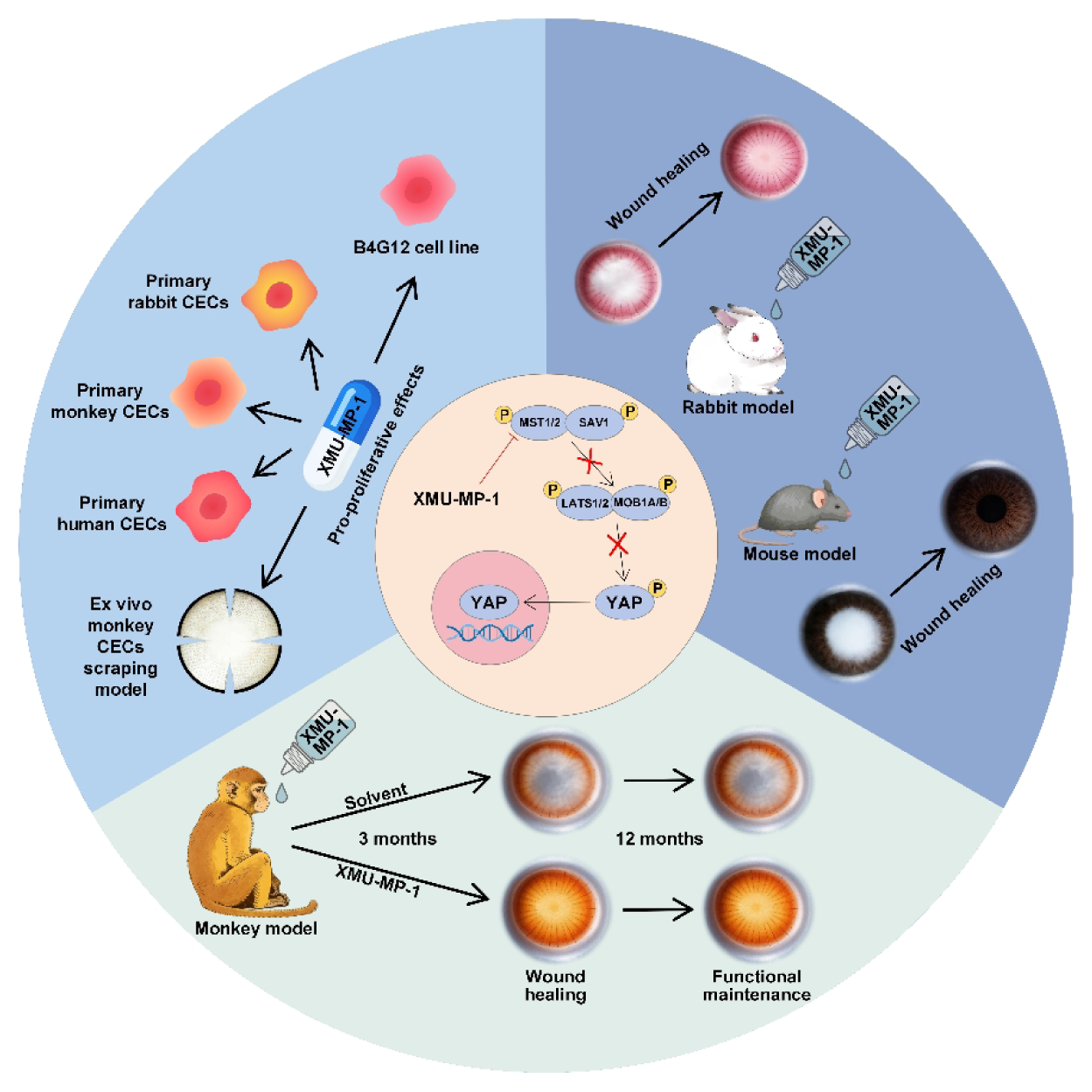
The cross-species pro-regenerative effect of XMU-MP-1. XMU-MP-1-induced activation of YAP promotes proliferation in B4G12 cells, primary rabbit CECs, monkey CECs and human CECs, and accelerates wound healing of *ex vivo* monkey corneal endothelium. XMU-MP-1 also demonstrates robust efficacy in promoting CECs regeneration in rabbit, mouse, and monkey.

For decades, how to regenerate human CECs has remained a huge challenge. Previous studies have demonstrated that nuclear translocation of YAP is associated with proliferation in primary culture of human CECs,^[32, 33]^ exogenous YAP could induce cell proliferation in contact-inhibited human CECs monolayers.^[34]^ However, the role of the Hippo pathway in CEC regeneration in vivo remains unknown. Our results derived in multi-species in vivo demonstrated that CECs regeneration is regulated by the Hippo pathway. In mouse corneal endothelial wounding model, EdU-positive cells were presented at the margin of injured region on day 3 post injury and expanded to nearly the entire damaged region on day 8 (Figure S9A, Supporting Information). Nevertheless, no EdU-positive cells were observed in the peripheral corneal endothelium (Figure S9A, Supporting Information) and the uninjured corneal endothelium (Figure S9B, Supporting Information), indicating the wound healing process initiates in the CECs at the wound margin, and the cell regeneration process is associated with the inactivation of Hippo pathway. Thus we speculate that the inability of human CECs to regenerate may due to evolutionary loss of the capacity to block the Hippo pathway following injury, a capacity which is retained in rodents.

Notably, YAP activation is also associated with endothelial-to-mesenchymal transition (EndMT),^[6]^ a pathological process during CEC wound healing characterized by upregulation of fibroblast biomarkers.^[35]^ Our previous study has found that rabbit CECs underwent a transient fibrotic EndMT following injury, and reversed back to an endothelial phenotype upon completion of wound healing.^[35]^ In this study, YAP activation has been demonstrated to promote cell cycle progression of CECs, and the CECs morphology could be restored to relatively normal state over time, concomitant with functional recovery of the corneal endothelium (Figure S10, Supporting Information). Therefore, EndMT may be also involved in the wound healing of CECs that regulated by YAP.

Although XMU-MP-1 has been shown to effectively promote corneal endothelial regeneration, potential side effects arising from YAP activation^[36]^ should be carefully considered. Our evaluation demonstrated that XMU-MP-1 exhibited ocular safety in mice following 4 weeks of topical administration (Figure S8, Supporting Information). Besides XMU-MP-1, TRULI and DMX-5804 are also pharmaceutical inhibitors for the Hippo pathway, which inhibit LATS1/2^[37]^ and non-canonical Hippo pathway MAPK4Ks^[38]^, respectively. In our study, topical administration of TRULI and DMX-5804 also showed effects in promoting CEC proliferation in vitro and in vivo (Figure S11, Supporting Information).

In the monkey experiments, we observed lower cell density and obviously irregular cell morphology in central corneal endothelium of the control group. Although one control cornea restored transparency 3 weeks post injury, which may be due to functional compensation of the surviving CECs, the CEC density of that control cornea did not restore to normal level. In comparison, XMU-MP-1 treatment restored both the cell density and morphology of central CECs to near normal levels, demonstrating the efficacy of XMU-MP-1 in promoting primate CECs regeneration. Based on the results obtained from the control group, it is reasonable to suspect that monkey CECs also possess a weak regenerative capacity in vivo, which is not identical to that of human. Given the difference between animal models and clinical conditions, future clinical study is necessary.

Our finding that a transient treatment with XMU-MP-1 can restore wounded corneal endothelium in non-human primates highlights a principle in regenerative biology: a properly timed signal modulation can initiate a complete program of tissue repair that extends the duration of the stimulus^[39–41]^. This contrasts with many chronic degenerative conditions that require prolonged pharmacological management. In the cornea endothelium, the injury itself may create a permissive microenvironment through changes in cell density, mechanical tension or extracellular matrix composition that allows a short pulse of YAP activation to drive not only proliferation but also the subsequent steps of polarity establishment, junctional reassembly, and functional maturation. Once a continuous monolayer of corneal endothelium is restored, contact inhibition and normalized cell-matrix signaling likely reactivate the Hippo pathway, thereby terminate the proliferative phase and preventing over-growth. This built-in feedback mechanism underscores the feasibility of targeting Hippo signaling in corneal endothelium.

## 4. Conclusion

In summary, we identified the Hippo signaling pathway as a regulator for CEC regeneration. Pharmaceutical inhibition of the Hippo pathway promotes corneal endothelial proliferation in multiple corneal endothelial wounding models across species, highlighting its potential for the treatment on corneal endothelial decompensation.

## 5. Methods

### Preparation of compounds

The solvent for XMU-MP-1 (MCE, HY-100526) and verteporfin (MCE, HY-B0146) consisted of 5% dimethyl sulfoxide (DMSO, MCE, HY-Y0320C), 20% polyethylene glycol 300 (PEG300, MCE, HY-Y0873) and 75% phosphate buffered solution (PBS, Solarbio, P1020).

### Animal

All animals, regardless of gender, were housed in Xiamen University Laboratory Animal Center and maintained under a 12-h light/dark cycle, free access to water and food. Animals used in this study were treated in accordance with the ARVO Statement for the Use of Animals in Ophthalmic and Vision Research.

### Animal examination

Corneal images were obtained using the slit-lamp microscope (TAKAGI, 700GL) and high-definition camera (Canon, EOS 7D). Anterior segment OCT images of rabbits and monkeys were obtained by swept-source OCT (TowardPi, Yalkaid YG-100K) and Small Animal Retina Imaging System (Optoprobe) was used for mice. A laser-scanning in vivo confocal microscope with a corneal imaging module (Heidelberg, HRT3-RCM) was used to observe the central CECs of monkeys, and the method of cell counting refers to our previous study.^[42]^

### Cynomolgus monkey corneal endothelial wound model

Central corneal endothelium damage was performed using transcorneal cryogenic method as previously described.^[43]^ The corneal endothelium of two cynomolgus monkeys was damaged under general anesthesia by transcorneal freezing with a 5-mm-diameter probe for 20 seconds. Before freezing, the copper probe was immersed in liquid nitrogen for 3 minutes to stabilize its temperature at approximately -196℃.

To evaluate the effect of XMU-MP-1 on CECs wound healing, XMU-MP-1 (50 μM) was topically applied to the right eye of each animal 4 times daily for 3 weeks. Solvent was applied to the lateral eye as a control. The experiments on two eyes were conducted separately to avoid affecting the monkeys in their daily activities.

### Rabbit corneal endothelial wound model

The corneal endothelium of New Zealand white rabbits was damaged under general anesthesia by transcorneal freezing with a 6-mm-diameter probe for 15 seconds. XMU-MP-1 (50 μM) or verteporfin (50 μM) was topically applied to rabbit eyes 4 times daily after corneal endothelial wounding. For the control group, solvent was administrated.

### Mouse corneal endothelial wound model

The UV-induced mouse corneal endothelial injury was performed as previously reported with modification.^[44]^ In brief, a UVB LED light source (Spectronics Corporation, EB-160C) with an emission peak of 312 nm light, 40 nm bandwidth and irradiance of 620 μW/cm^2^ was focused to illuminate a 2-mm-diameter spot onto the mouse cornea for 7 min with the total irradiance energy of 260.4 J/cm^2^. After irradiation, 2 μl of XMU-MP-1 (1 μM) or verteporfin (50 μM) was topically applied to mouse eyes 4 times daily, and solvent was applied to the control group.

### AAV injection in mouse

The AAV was manufactured by Hanbio Biotechnology (Shanghai, China). To achieve endothelial-selective gene knockdown, sequence encoding shYap1 or negative control sequence was constructed under Tie1 promoter and expressed in AAV (Table S1). The virus titer was 1.6×10^12^ μg/ml.

Then, 8-week-old C57BL/6 mice were anesthetized and anterior chamber was injected with AAV Control-ZsGreen or AAV Yap1 shRNA-ZsGreen (3 μl per eye) using a 34-gauge needle. ZsGreen expression was detected in vivo using stereo fluorescence microscope (Leica, M165FC) 4 weeks after injection, then the mice were used to generate corneal endothelial wound model.

### Evaluation of corneal transparency

The cornea was observed under slit-lamp microscope and the corneal transparency was evaluated with scores ranged from 0 to 4 as previously reported: 0, a normal cornea; 0.5, a slight haze visible only under a slit-lamp microscope; 1, a mild haze; 2, a moderate haze with a visible iris; 3, a severe haze with an invisible iris; and 4, a severe haze with corneal ulceration.^[45]^

### Ex vivo monkey corneal endothelial scraping model

Two monkey corneas were divided into 12 equal pieces. After culturing for 24 hours with the endothelial surface facing upward, CECs were scraped off within a 2 mm diameter area using a soft rubber rod. Trypan blue staining was used to visualize areas of endothelial defect. The corneal explants were cultured in Dulbecco’s Modified Eagle Medium/Nutrient Mixture F-12 (DMEM/F12, Gibco, 11320033) supplemented with 5% fetal bovine serum (FBS, ABW, AB-FBS-0050), 0.5% DMSO, 2 ng/mL mouse EGF (Gibco, PMG8041), 1% Insulin-Transferrin-Selenium-Ethanolamine (ITS-X, Gibco, A4000046401), 0.5 μg/mL hydrocortisone (Yeasen, 54576ES03) and 1% penicillin/streptomycin (P/S, Gibco, 15140-122), with culture medium changed every other day.

### Isolation and primary culture of monkey CECs

Monkey CECs were isolated through Descemet’s stripping and digestion with 1 mg/mL collagenase A (Merck, 10103578001) for 6 h at 37 ℃. After 5 min in TrypLE (Thermo Fisher Scientific, 12605010) at 37℃, CECs were pelleted at 1200 rpm for 5 min and plated on fibronectin coating mix (AthenaES, 0407) treated wells. The cells were cultured in Dulbecco’s Modified Eagle Medium (DMEM, Gibco, 11965092) with 10% FBS at 37℃ in 5% CO_2_.

### Isolation and primary culture of human CECs and rabbit CECs

Human tissue was handled according to the Declaration of Helsinki. Corneoscleral tissues from human donor eyes were obtained from the Eye Bank of Xiamen Eye Center of Xiamen University immediately after the central corneal button had been used for corneal transplantation. Human CECs and rabbit CECs were isolated using our previous method,^[46]^ and the cells were cultured in DMEM/F12 supplemented with 5% FBS, 0.5% DMSO, 2 ng/mL mouse EGF, 1% ITS-X, 0.5 μg/mL hydrocortisone and 1% P/S.

### Transfection of rabbit CECs and B4G12 with siRNA

YAP expression was knocked down in rabbit CECs and B4G12 cell line by transfection with the YAP siRNA (Sangon Biotech) using RNATransMate (Sangon Biotech, E607402) following the manufacturer’s instruction, with negative control (NC) siRNA as control. The sequences of the siRNAs are listed in Table S2.

### In vitro and in vivo EdU labeling

For *in vitro* EdU labeling, EdU (APExBIO, B8337) was supplemented in the culture medium for a final concentration of 20 mg/ml. Cells were collected after 8 hours labeling.

For *in vivo* EdU labeling, mice were given an intraperitoneal injection with 50 mg/kg of EdU in sterile PBS for 8 consecutive days. After that, the animals were sacrificed and the corneas were collected for cryosection. EdU signal in corneal endothelium was detected using the Click-iT EdU Alexa Fluor 647 imaging kit (Invitrogen, C10640) according to the manufacturer’ s instruction.

### CCK-8 assay

The CCK-8 solution (Vazyme, A311-01) was added to the culture medium, and the absorbance value (450 nm) for each well was determined 2 hours later using a microplate reader (Thermo, Multiskan GO).

### Western blot analysis

Western blot analysis was performed as previously described.^[47]^ The equal amount of protein was resolved by SDS-PAGE and immunoblotted with primary antibodies for YAP1 (Abclonal, A19134), phospho-YAP1-S127 (Abcam, ab186735), LATS1/2 (Proteintech, 17049-1-AP), phospho-LATS1/2 (Thr1079) (Proteintech, 28998-1-AP), MST1/2 (Proteintech, 22245-1-AP), phospho-MST1 (Thr183)/MST2 (Thr180) (Proteintech, 80093-1-RR), MOB1 (Proteintech, 11669-1-AP), phospho-MOB1 (Thr12) (Proteintech, 29027-1-AP) and β-actin (Abcam, 8226). Primary antibodies were then detected with HRP-conjugated goat anti-rabbit antibody (Abclonal, AS014), and the band intensity was semiquantified by densitometry using ImageJ software and normalized by β-actin levels.

### Quantitative real-time polymerase chain reaction (qRT-PCR)

Total RNA was extracted using Total RNA Extraction Reagent (Abclonal, RK30129), and cDNA was made by using the RT Master Mix for qPCR (Abclonal, RK20433). Real-time qPCR was performed with the SYBR Green qPCR Mix with UDG (Abclonal, RK21219), and the primer sequences are listed in Table S3. The mixture was held at 37℃ for 2 min and then heated to 95℃ for 3 min, cycled 40 times (95℃ for 5 s, 60℃ for 30 s). Melting curves were generated by increasing the temperature from 55℃ to 95℃ in 0.5℃ increments at 10 s intervals and then visually to ensure that a single peak was present for each primer. Threshold amplification values (Ct) were assigned by the LightCycler 96 analysis software (Roche).

### Immunofluorescence staining on cells and corneal whole mount

Cells or corneas were fixed with 4% paraformaldehyde for 20 min and incubated with 2% bovine serum albumin for 60 min to reduce background noise. The cells or corneas were then labeled with primary rabbit antibodies for YAP1 (Abclonal, A19134), Ki67 (Abcam, ab16667), ZO-1 (Invitrogen, 61-7300) or Na^+^-K^+^ATPase (Abcam, ab76020) overnight at 4℃. After 3 washes with PBS, the samples were incubated with goat anti-rabbit lgG (Abclonal, AS053) for 120 min. The nuclei were stained with DAPI (Beyotime, C1005) for 10 min. The samples were observed under a laser scanning confocal microscopy (ZEISS, LSM880).

### Alizarin Red staining

Mouse or rabbit corneas were incubated with Alizarin Red S solution (Solarbio, G1452) for 5 min. After that, the corneas were fixed with 4% paraformaldehyde for 20 min and cut into flat mount. Images were captured with a microscope (Zeiss, Axio Lab.A1).

### Statistics

Statistical analysis was performed using GraphPad Prism software 9.0. Data were presented as mean±SEM and analyzed by unpaired Student’s *t* test for two groups and ANOVA for more than two groups. P < 0.05 was considered statistical significant.

### Ethics Approval Statement

The animal experiments were performed according to the protocol approved by the Laboratory Animal Management and Ethics Committee of Xiamen University (Approval No. XMULAC20240087).

## Supporting Information

Supporting Information is available from the Wiley Online Library or from the author.

## Acknowledgement

We thank Jingru Huang and Xiang You from Central Lab, School of Medicine, Xiamen University for technical support in confocal imaging. We thank Haiping Zheng from Central Lab, School of Medicine, Xiamen University for technical support in cell cycle analysis.

## Conflict of interest

The authors have declared that no conflict of interest exists.

## Author contributions

J.Z., Y.G., H.Z. and W.L. designed the overall project and wrote the manuscript, J.Z., Y.G., H.Z., W.H., Y.W., M.Z., X.H., L.Z., J.Y. S.C. H.H. and R.Z. performed the experiments and data analysis. H.W. collected human corneoscleral tissues. Z.L., A.Q, C.C, and W.L. revised the manuscript. J.Z., Y.G., and H.Z. contributed equally to this work, thus should be considered as co-first authors. The order of co–first authors was decided by discussions among the 3 first authors and the 2 corresponding author. All authors have reviewed and approved the manuscript.

## Data Availability Statement

The data supporting the findings of this study are available in the Supporting Data Values file. The raw data used in this study are available from the corresponding author upon reasonable request.

## Notes

**Funding:** This study was supported in part by the National Natural Science Foundation of China (82471047, 81970773, 82401223), China Post-doctoral science foundation (2024M751351, 2023M741610, BX20240154), Natural Science Foundation of Hunan Province, China (2025JJ90130), Natural Science Foundation of Hunan Province, China (2024JJ6403).

### Competing Interest Statement

The authors have declared no competing interest.

### Summary of Updates

The sequencing-related data removed from the Figure 1; Figure 5 and Figure 6 added; Figure 4 revised; 2 authors added; Supplemental files updated; Layout format updated.

## References

[1] K.M. Meek, C. Knupp, Corneal structure and transparency, Prog Retin Eye Res, 49 (2015) 1–16.

[2] G. Suanno, V.G. Genna, E. Maurizi, A. Abu Dieh, M. Griffith, G. Ferrari, Cell therapy in the cornea: The emerging role of microenvironment, Prog Retin Eye Res, 102 (2024).

[3] W.M. Bourne, Biology of the corneal endothelium in health and disease, Eye, 17 (2003) 912–918.

[4] K. Gupta, S.X. Deng, Corneal Endothelial Decompensation, Klin Monatsbl Augenh, 237 (2020) 745–753.

[5] Z.G. He, N. Campolmi, P. Gain, M.H. Binh, J.M. Dumollard, S. Duband, M. Peoc’h, S. Piselli, O. Garraud, G. Thuret, Revisited Microanatomy of the Corneal Endothelial Periphery: New Evidence for Continuous Centripetal Migration of Endothelial Cells in Humans, Stem Cells, 30 (2012) 2523–2534.

[6] X. Zhang, H. Liu, C. Wan, Y. Li, C. Ren, J. Lu, Y. Liu, Y. Yang, Verteporfin combined with ROCK inhibitor promotes the restoration of corneal endothelial cell dysfunction in rats, Biochem Pharmacol, 231 (2025) 116641.

[7] A. Viberg, B. Samolov, B. Bystroem, Descemet Stripping Automated Endothelial Keratoplasty versus Descemet Membrane Endothelial Keratoplasty for Fuchs Endothelial Corneal Dystrophy, Ophthalmology, 130 (2023) 1248–1257.

[8] M.O. Price, P. Calhoun, C. Kollman, F.W. Price, J.H. Lass, Descemet Stripping Endothelial Keratoplasty, Ophthalmology, 123 (2016) 1421–1427.

[9] S. Kinoshita, N. Koizumi, M. Ueno, N. Okumura, K. Imai, H. Tanaka, Y. Yamamoto, T. Nakamura, T. Inatomi, J. Bush, M. Toda, M. Hagiya, I. Yokota, S. Teramukai, C. Sotozono, J. Hamuro, Injection of Cultured Cells with a ROCK Inhibitor for Bullous Keratopathy, New Engl J Med, 378 (2018) 995–1003.

[10] K. Numa, K. Imai, M. Ueno, K. Kitazawa, H. Tanaka, J.D. Bush, S. Teramukai, N. Okumura, N. Koizumi, J. Hamuro, C. Sotozono, S. Kinoshita, Five-Year Follow-up of First 11 Patients Undergoing Injection of Cultured Corneal Endothelial Cells for Corneal Endothelial Failure, Ophthalmology, 128 (2021) 504–514.

[11] Z.Y. Li, H.Y. Duan, Y.N. Jia, C. Zhao, W.J. Li, X. Wang, Y.J. Gong, C.X. Dong, B.C. Ma, S.Q. Dou, B. Zhang, D.F. Li, Y.H. Cao, L.X. Xie, Q.J. Zhou, W.Y. Shi, Long-term corneal recovery by simultaneous delivery of hPSC-derived corneal endothelial precursors and nicotinamide, J Clin Invest, 132 (2022).

[12] N. Okumura, S. Nakano, E.P. Kay, R. Numata, A. Ota, Y. Sowa, T. Sakai, M. Ueno, S. Kinoshita, N. Koizumi, Involvement of Cyclin D and p27 in Cell Proliferation Mediated by ROCK Inhibitors Y-27632 and Y-39983 During Corneal Endothelium Wound Healing, Invest Ophth Vis Sci, 55 (2014) 318–329.

[13] L.C. Meekins, N. Rosado-Adames, R. Maddala, J.J. Zhao, P.V. Rao, N.A. Afshari, Corneal Endothelial Cell Migration and Proliferation Enhanced by Rho Kinase (ROCK) Inhibitors in In Vitro and In Vivo Models, Invest Ophth Vis Sci, 57 (2016) 6731–6738.

[14] N. Okumura, Y. Okazaki, R. Inoue, K. Kakutani, S. Nakano, S. Kinoshita, N. Koizumi, Effect of the Rho-Associated Kinase Inhibitor Eye Drop (Ripasudil) on Corneal Endothelial Wound Healing, Invest Ophth Vis Sci, 57 (2016) 1284–1292.

[15] H. Miyagi, S. Kim, J. Li, C.J. Murphy, S.M. Thomasy, Topical Rho-Associated Kinase Inhibitor, Y27632, Accelerates Corneal Endothelial Regeneration in a Canine Cryoinjury Model, Cornea, 38 (2019) 352–359.

[16] N. Koizumi, N. Okumura, M. Ueno, H. Nakagawa, J. Hamuro, S. Kinoshita, Rho-Associated Kinase Inhibitor Eye Drop Treatment as a Possible Medical Treatment for Fuchs Corneal Dystrophy, Cornea, 32 (2013) 1167–1170.

[17] G. Moloney, C. Petsoglou, M. Ball, Y. Kerdraon, R. Höllhumer, N. Spiteri, S. Beheregaray, J. Hampson, M. D’Souza, R.N. Devasahayam, Descemetorhexis Without Grafting for Fuchs Endothelial Dystrophy-Supplementation With Topical Ripasudil, Cornea, 36 (2017) 642–648.

[18] N. Okumura, S. Kinoshita, N. Koizumi, Application of Rho Kinase Inhibitors for the Treatment of Corneal Endothelial Diseases, J Ophthalmol, 2017 (2017).

[19] W.D. Pei, J. Chen, W.Y. Wu, W. Wei, Y. Yu, Y. Feng, Comparison of the rabbit and human corneal endothelial proteomes regarding proliferative capacity, Exp Eye Res, 209 (2021).

[20] K. Yoshida, S. Kase, K. Nakayama, H. Nagahama, T. Harada, H. Ikeda, C. Harada, J. Imaki, K. Ohgami, K. Shiratori, I.B. Ilieva, S. Ohno, S. Nishi, K.I. Nakayama, Involvement of p27 in the proliferation of the developing corneal endothelium, Invest Ophth Vis Sci, 45 (2004) 2163–2167.

[21] S.P. Hui, D.Z. Sheng, K. Sugimoto, A. Gonzalez-Rajal, S. Nakagawa, D. Hesselson, K. Kikuchi, Zebrafish Regulatory T Cells Mediate Organ-Specific Regenerative Programs, Dev Cell, 43 (2017) 659–+.

[22] P. Lyu, M. Iribarne, D. Serjanov, Y.J. Zhai, T. Hoang, L.J. Campbell, P. Boyd, I. Palazzo, M. Nagashima, N.J. Silva, P.F. Hitchcock, J. Qian, D.R. Hyde, S. Blackshaw, Common and divergent gene regulatory networks control injury-induced and developmental neurogenesis in zebrafish retina, Nat Commun, 14 (2023).

[23] L. Celotto, F. Rost, A. Machate, J. Bläsche, A. Dahl, A. Weber, S. Hans, M. Brand, Single-cell RNA sequencing unravels the transcriptional network underlying zebrafish retina regeneration, Elife, 12 (2023).

[24] A. Elbediwy, Z.I. Vincent-Mistiaen, B. Spencer-Dene, R.K. Stone, S. Boeing, S.K. Wculek, J. Cordero, E.H. Tan, R. Ridgway, V.G. Brunton, E. Sahai, H. Gerhardt, A. Behrens, I. Malanchi, O.J. Sansom, B.J. Thompson, Integrin signalling regulates YAP and TAZ to control skin homeostasis, Development, 143 (2016) 1674–1687.

[25] Z. Liu, H. Wu, K. Jiang, Y. Wang, W. Zhang, Q. Chu, J. Li, H. Huang, T. Cai, H. Ji, C. Yang, N. Tang, MAPK-Mediated YAP Activation Controls Mechanical-Tension-Induced Pulmonary Alveolar Regeneration, Cell Rep, 16 (2016) 1810–1819.

[26] M. Xin, Y. Kim, L.B. Sutherland, M. Murakami, X.X. Qi, J. McAnally, E.R. Porrello, A.I. Mahmoud, W. Tan, J.M. Shelton, J.A. Richardson, H.A. Sadek, R. Bassel-Duby, E.N. Olson, Hippo pathway effector Yap promotes cardiac regeneration, P Natl Acad Sci USA, 110 (2013) 13839–13844.

[27] L. Lu, Y. Li, S.M. Kim, W. Bossuyt, P. Liu, Q. Qiu, Y. Wang, G. Halder, M.J. Finegold, J.S. Lee, R.L. Johnson, Hippo signaling is a potent in vivo growth and tumor suppressor pathway in the mammalian liver, Proc Natl Acad Sci U S A, 107 (2010) 1437–1442.

[28] F.X. Yu, K.L. Guan, The Hippo pathway: regulators and regulations, Gene Dev, 27 (2013) 355–371.

[29] J.B. Huang, S. Wu, J. Barrera, K. Matthews, D.J. Pan, The Hippo signaling pathway coordinately regulates cell proliferation and apoptosis by inactivating Yorkie, the homolog of YAP, Cell, 122 (2005) 421–434.

[30] A. Perra, M.A. Kowalik, E. Ghiso, G.M. Ledda-Columbano, L. Di Tommaso, M.M. Angioni, C. Raschioni, E. Testore, M. Roncalli, S. Giordano, A. Columbano, YAP activation is an early event and a potential therapeutic target in liver cancer development, J Hepatol, 61 (2014) 1088–1096.

[31] F.Q. Fan, Z.X. He, L.L. Kong, Q.H. Chen, Q. Yuan, S.H. Zhang, J.J. Ye, H. Liu, X.F. Sun, J. Geng, L.Z. Yuan, L.X. Hong, C. Xiao, W.J. Zhang, X.H. Sun, Y.Z. Li, P. Wang, L.H. Huang, X.R. Wu, Z.L. Ji, Q. Wu, N.S. Xia, N.S. Gray, L.F. Chen, C.H. Yun, X.M. Deng, D.W. Zhou, Pharmacological targeting of kinases MST1 and MST2 augments tissue repair and regeneration, Sci Transl Med, 8 (2016).

[32] Y.T. Zhu, H.C. Chen, S.Y. Chen, S.C.G. Tseng, Nuclear p120 catenin unlocks mitotic block of contact-inhibited human corneal endothelial monolayers without disrupting adherent junctions, J Cell Sci, 125 (2012) 3636–3648.

[33] C. Zhao, Q.J. Zhou, H.Y. Duan, X. Wang, Y.N. Jia, Y.J. Gong, W.J. Li, C.X. Dong, Z.Y. Li, W.Y. Shi, Laminin 511 Precoating Promotes the Functional Recovery of Transplanted Corneal Endothelial Cells, Tissue Eng Pt A, 26 (2020) 1158–1168.

[34] Y.J. Hsueh, H.C. Chen, S.E. Wu, T.K. Wang, J.K. Chen, D.H.K. Ma, Lysophosphatidic acid induces YAP-promoted proliferation of human corneal endothelial cells via PI3K and ROCK pathways, Mol Ther-Meth Clin D, 2 (2015).

[35] J. Chen, Z. Li, L. Zhang, S. Ou, Y. Wang, X. He, D. Zou, C. Jia, Q. Hu, S. Yang, X. Li, J. Li, J. Wang, H. Sun, Y. Chen, Y.T. Zhu, S.C.G. Tseng, Z. Liu, W. Li, Descemet’s Membrane Supports Corneal Endothelial Cell Regeneration in Rabbits, Sci Rep, 7 (2017) 6983.

[36] I.M. Moya, G. Halder, Hippo-YAP/TAZ signalling in organ regeneration and regenerative medicine, Nat Rev Mol Cell Bio, 20 (2019) 211–226.

[37] N. Kastan, K. Gnedeva, T. Alisch, A.A. Petelski, D.J. Huggins, J. Chiaravalli, A. Aharanov, A. Shakked, E. Tzahor, A. Nagiel, N. Segil, A.J. Hudspeth, Small-molecule inhibition of Lats kinases may promote Yap-dependent proliferation in postmitotic mammalian tissues, Nat Commun, 12 (2021).

[38] L.R. Fiedler, K. Chapman, M. Xie, E. Maifoshie, M. Jenkins, P.A. Golforoush, M. Bellahcene, M. Noseda, D. Faust, A. Jarvis, G. Newton, M.A. Paiva, M. Harada, D.J. Stuckey, W.H. Song, J. Habib, P. Narasimham, R. Aqil, D. Sanmugalingam, R. Yan, L. Pavanello, M. Sano, S.C. Wang, R.D. Sampson, S. Kanayaganam, G.E. Taffet, L.H. Michae, M.L. Entman, T.H. Tan, S.E. Harding, C.M.R. Low, C. Tralau-Stewart, T. Perrior, M.D. Schneider, MAP4K4 Inhibition Promotes Survival of Human Stem Cell-Derived Cardiomyocytes and Reduces Infarct Size, Cell Stem Cell, 24 (2019) 579–+.

[39] A. Ocampo, P. Reddy, P. Martinez-Redondo, A. Platero-Luengo, F. Hatanaka, T. Hishida, M. Li, D. Lam, M. Kurita, E. Beyret, T. Araoka, E. Vazquez-Ferrer, D. Donoso, J.L. Roman, J. Xu, C. Rodriguez Esteban, G. Nuñez, E. Nuñez Delicado, J.M. Campistol, I. Guillen, P. Guillen, J.C. Izpisua Belmonte, In Vivo Amelioration of Age-Associated Hallmarks by Partial Reprogramming, Cell, 167 (2016) 1719–1733.e1712.

[40] D. Yimlamai, C. Christodoulou, G.G. Galli, K. Yanger, B. Pepe-Mooney, B. Gurung, K. Shrestha, P. Cahan, B.Z. Stanger, F.D. Camargo, Hippo pathway activity influences liver cell fate, Cell, 157 (2014) 1324–1338.

[41] M. Xin, Y. Kim, L.B. Sutherland, M. Murakami, X. Qi, J. McAnally, E.R. Porrello, A.I. Mahmoud, W. Tan, J.M. Shelton, J.A. Richardson, H.A. Sadek, R. Bassel-Duby, E.N. Olson, Hippo pathway effector Yap promotes cardiac regeneration, Proc Natl Acad Sci U S A, 110 (2013) 13839–13844.

[42] J.H. Bu, J.W. Yu, Y. Wu, X.X. Cai, K.C. Li, L.Y. Tang, N. Jiang, M.V. Jeyalatha, M.J. Zhang, H.M. Sun, H. He, A.J. Quantock, Y.X. Chen, Z.G. Liu, W. Li, Hyperlipidemia Affects Tight Junctions and Pump Function in the Corneal Endothelium, Am J Pathol, 190 (2020) 563–576.

[43] N. Okumura, N. Koizumi, E.P. Kay, M. Ueno, Y. Sakamoto, S. Nakamura, J. Hamuro, S. Kinoshita, The ROCK Inhibitor Eye Drop Accelerates Corneal Endothelium Wound Healing, Invest Ophth Vis Sci, 54 (2013) 2493–2502.

[44] C.L. Liu, T. Miyajima, G. Melangath, T. Miyai, S. Vasanth, N. Deshpande, V. Kumar, S.O. Tone, R. Gupta, S. Zhu, D. Vojnovic, Y.M. Chen, E.G. Rogan, B. Mondal, M. Zahid, U.V. Jurkunas, Ultraviolet A light induces DNA damage and estrogen-DNA adducts in Fuchs endothelial corneal dystrophy causing females to be more affected, P Natl Acad Sci USA, 117 (2020) 573–583.

[45] C. Zhao, W. Li, H. Duan, Z. Li, Y. Jia, S. Zhang, X. Wang, Q. Zhou, W. Shi, NAD(+) precursors protect corneal endothelial cells from UVB-induced apoptosis, Am J Physiol Cell Physiol, 318 (2020) C796–c805.

[46] W. Li, A.L. Sabater, Y.T. Chen, Y. Hayashida, S.Y. Chen, H. He, S.C.G. Tseng, A novel method of isolation, preservation, and expansion of human corneal endothelial cells, Invest Ophth Vis Sci, 48 (2007) 614–620.

[47] S. Li, L. Tang, J. Zhou, S. Anchouche, D. Li, Y. Yang, Z. Liu, J. Wu, J. Hu, Y. Zhou, J. Yin, Z. Liu, W. Li, Sleep deprivation induces corneal epithelial progenitor cell over-expansion through disruption of redox homeostasis in the tear film, Stem Cell Reports, 17 (2022) 1105–1119.

