## Supplementary methods and figures for "The Hippo Signaling Pathway Regulates Corneal Endothelial Regeneration"

W. Huang

TowardPi Medical Technology Beijing; Beijing, China.

A. Quantock

School of Optometry and Vision Sciences, Cardiff University, Cardiff, UK.

**Supplementary Methods**

**B4G12 cell line**

B4G12 cell line was purchase from OTWO Biotech (HTX1856) and tested for mycoplasma contamination and was authenticated May 2025 by STR sequencing. The culture medium was composed of DMEM, 10% FBS and PS.

**Cell cycle analysis**

Rabbit CECs were harvested and fixed in 70% ethanol for 2 hours at 4°C and then stained with Cell Cycle Analysis Kit (Beyotime, C1052) according to the manufacturer’s protocol. Then rabbit CECs were examined with a flow cytometer (Beckman, CytoFLEX), and counts of G1, S, and G2/M phase cells were determined with the ModFIT software. Each experiment was repeated three independent times.

**In vitro primary monkey CECs scraping model**

Primary monkey CECs were cultured in 6-well plates until complete confluence, then the cells in the center of the well were scraped off with a diameter of 1 cm using sterile rubber rods.

**Assessment of side effect of XMU-MP-1 treatment**

Mice eyes were topically applied with 1 μM of XMU-MP-1 4 times daily for 4 weeks, with solvent as control. Fluorescein sodium staining images were photographed with Cobalt blue filter. Tear production was tested by wetted phenol thread lengths without topical anesthesia. The intraocular pressure (IOP) was measured by a tonometer (Tonovet). Axial length of eyes was measured by swept-source OCT (TowardPi, Yalkaid YG-100K), and the refraction was tested by Photorefractor (Striatech). OCT images of mice retina were obtained by SKY-lab MAX (Micro Clear) under general anesthesia, and the fundus fluorescein angiography (FFA) was conducted using Small Animal Retina Imaging System (Optoprobe) after intraperitoneal injection of fluorescein sodium. The eyeballs and peripheral conjunctival tissue, Meibomian glands and lacrimal glands were carefully collected after the animals were sacrificed. Hematoxylin & eosin (HE) staining was performed on the cryosections of the tissues.

**Liquid chromatography-tandem mass spectrometry (LC-MS) analysis of XMU-MP-1**

LC separation was performed using a Nexera X2 LC-30AD ultra-HPLC system. The mobile phase flow rate was 300 μL/min, and the mobile phases were 0.1% formic acid in water (mobile phase A) and 0.1% formic acid in acetonitrile (mobile phase B). The LC gradient conditions were as follows: 10%-90% B for 0−4 min, 90% B for 4−6 min, 90%-10% B for 6-7 min, 10% B for 7-10 min.

Mass detection was carried out using positive ionization in multiple reaction monitoring mode (MRM). The nebulizer gas (GS1) was nitrogen at 55 psi, and the auxiliary gas (GS2) was nitrogen maintained at 55 psi. The collision (CAD) and curtain gas (CUR) values were set to 35 psi, respectively. The ion spray voltage was set to 5500 V in positive mode, and the heater temperature was 550℃. The system control and data analysis was performed by AB Sciex Analyst® software. The detected ion pairs were 417 > 220* (quantitative ion pair) and 417 > 390* (qualitative ion pair).

Analyst 1.6.3 software was used to extract chromatographic peak areas and retention times. According to the retention time and peak type information of the standard, the qualitative and quantitative analysis of all samples was carried out. The peak area of each chromatographic peak was substituted for the relative content of the corresponding substance, and then substituted into the linear equation and calculation formula, and finally the qualitative and quantitative analysis results of XMU-MP-1 in all samples were obtained.

For preparation of samples, the rabbit corneal endothelium was separated after topical application of XMU-MP-1 (50μM) for 3 days. The tissue was homogenized in pre-cooled methanol/acetonitrile/water (2/2/1) and sonicated for 30 min. The supernatant was then centrifuged and subjected to LC-MS/MS analysis.

**Supplementary Figures and Legends**

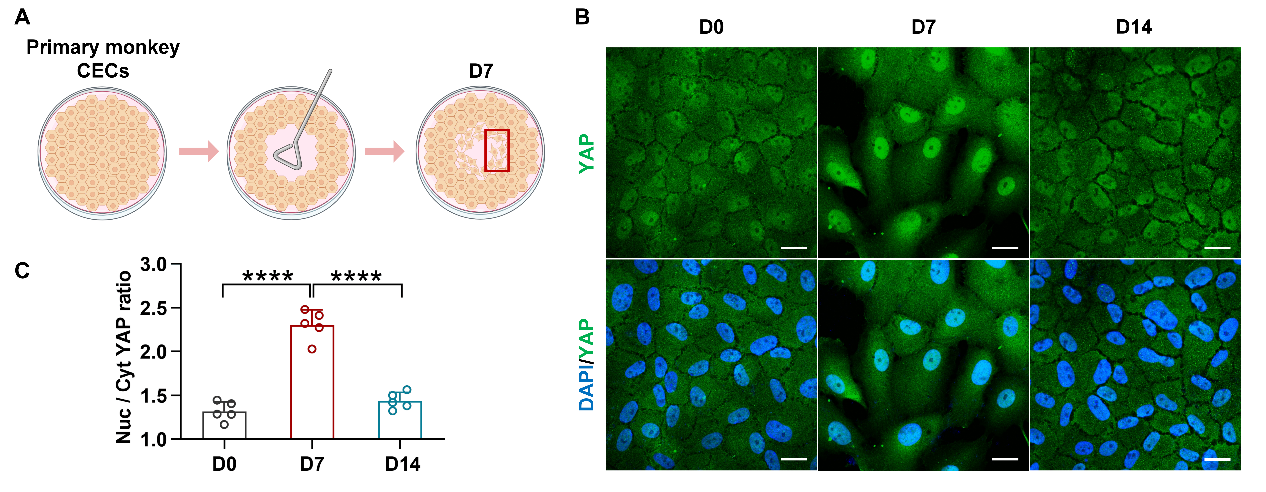

**Figure S1. Nuclear translocation of YAP in *in vitro* monkey CEC scraping model.** (A) The diagram shows the *in vitro* monkey CEC scraping model. (B) Representative immunofluorescence images of YAP in primary monkey CECs. (C) The ratio of nuclear YAP to cytoplasm YAP in primary monkey CECs, five cells were randomly selected for measurement and averaged in each sample (n = 3). ****P < 0.0001, error bars, mean ± SEM. Scale bars represent 15 µm in (B). Nuc, nucleus; Cyt, cytoplasm.

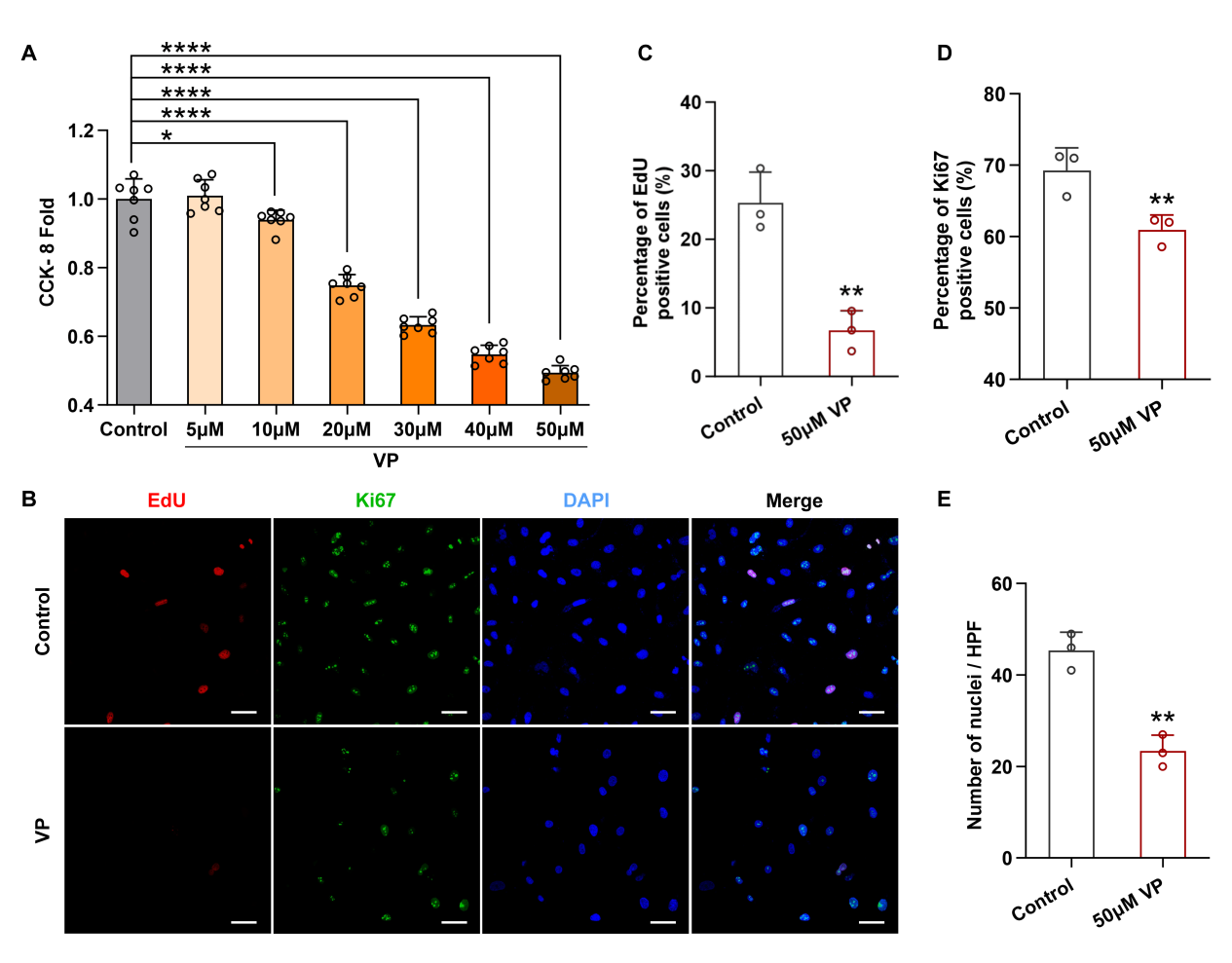

**Figure S2. Verteporfin suppresses proliferation of B4G12 cell line.** (A) CCK-8 assay of B4G12 cell line treated with Verteporfin (Fold of control, n=7). (B) Representative EdU staining and immunofluorescence images of Ki67 in B4G12. (C to E) Percentages of EdU (C) and Ki67 (D) positive cells and nuclei counting (E) in (B) (n=3). Three visual fields were randomly selected in each sample for statistical processing. *P < 0.05, **P < 0.01, ***P < 0.001, ****P < 0.0001, error bars, mean ± SEM. Scale bars represent 50 µm. VP, verteporfin.

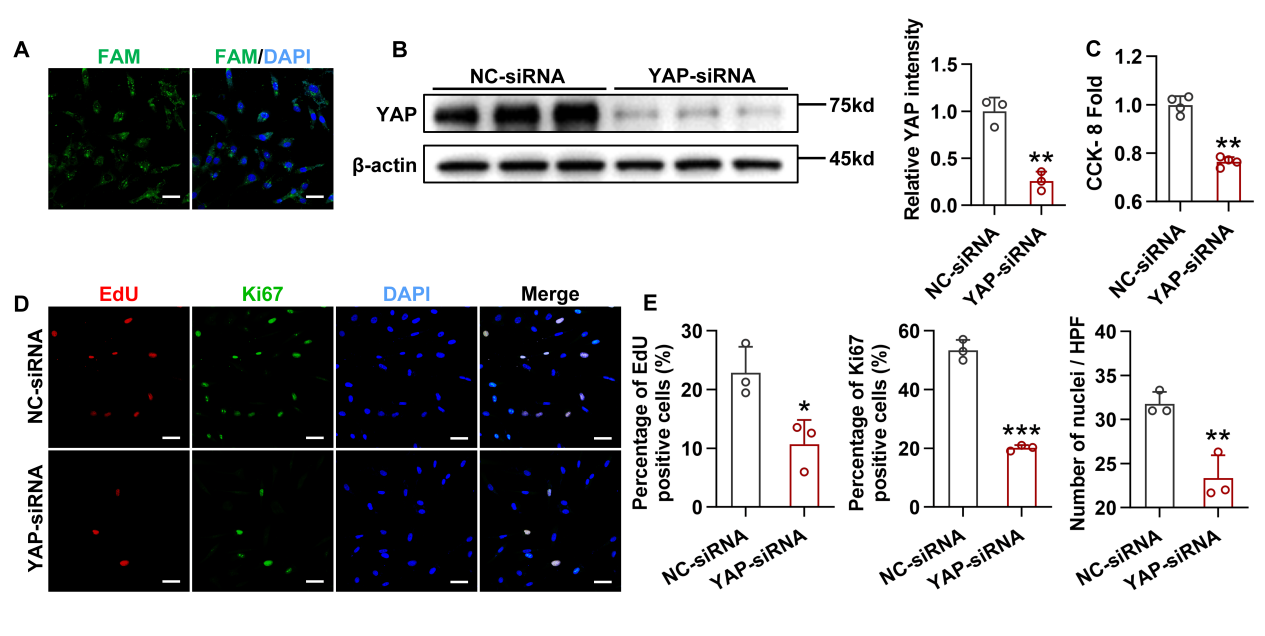

**Figure S3. siRNA-mediated knockdown of *Yap1* inhibits proliferation of B4G12 cell line.** (A) FAM expression in B4G12 cells transfected with NC siRNA or YAP siRNA, demonstrating the success of transfection. (B) YAP expression and densitometry quantification in the B4G12 cells after transfection (n = 3). (C) CCK-8 assay of B4G12 cells (Fold of control, n = 4). (D) Representative EdU and Ki67 staining in B4G12 cells. (E) Percentages of EdU and Ki67 positive cells and nuclei counting in (D) (n=3). Three visual fields were randomly selected in each sample for statistical processing. *P < 0.05, **P < 0.01, ***P < 0.001, error bars, mean ± SEM. Scale bars represent 50 µm. NC, negative control.

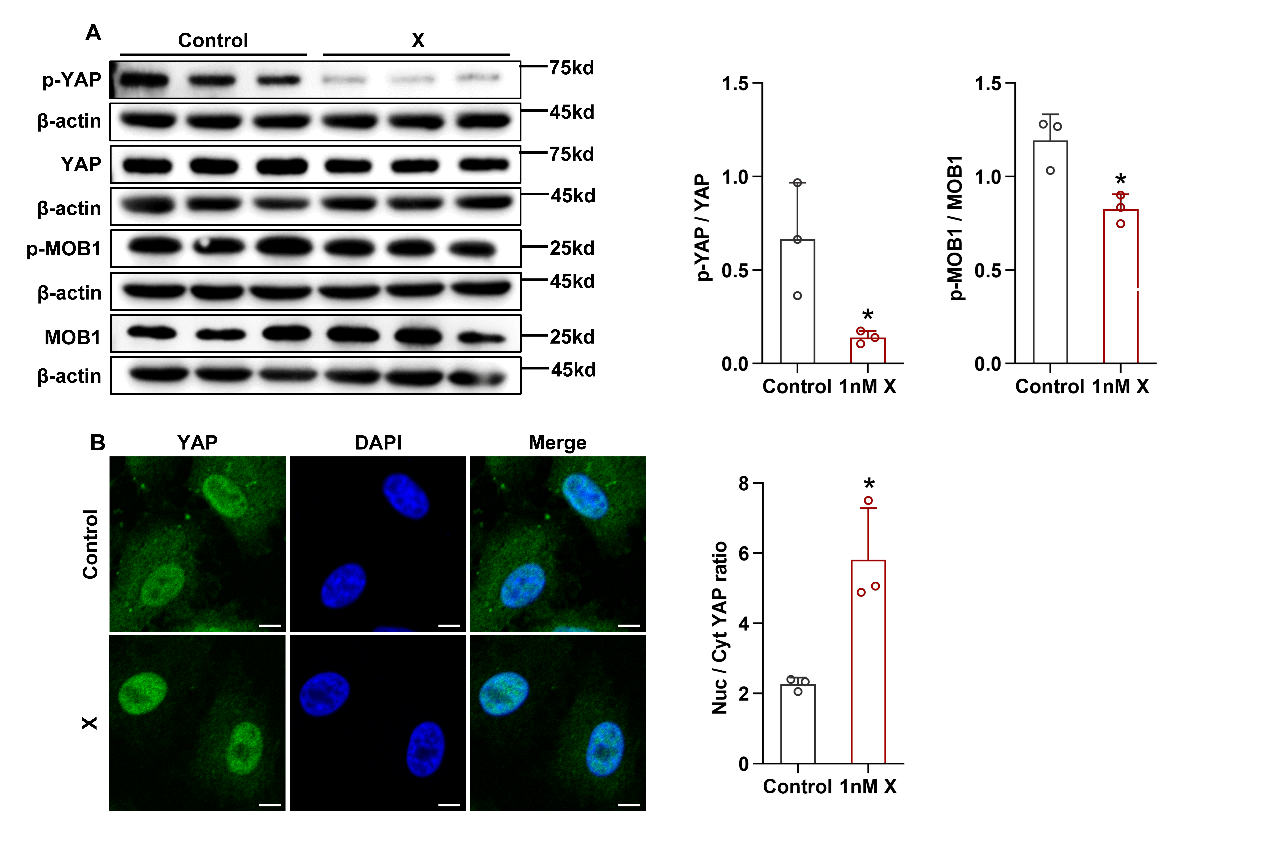

**Figure S4. XMU-MP-1 inhibits YAP phosphorylation and enhances YAP nuclear translocation.** (A) Western blot analysis of p-YAP, YAP, p-MOB1 and MOB1 in primary rabbit CECs and the ratios of p-YAP to YAP and p-MOB1 to MOB1 (n = 3). (B) Representative immunofluorescence images of YAP in primary rabbit CECs and the ratio of nuclear YAP to cytoplasm YAP (n = 3). *P < 0.05, error bars, mean ± SEM. Scale bars represent 6 µm. X, XMU-MP-1.

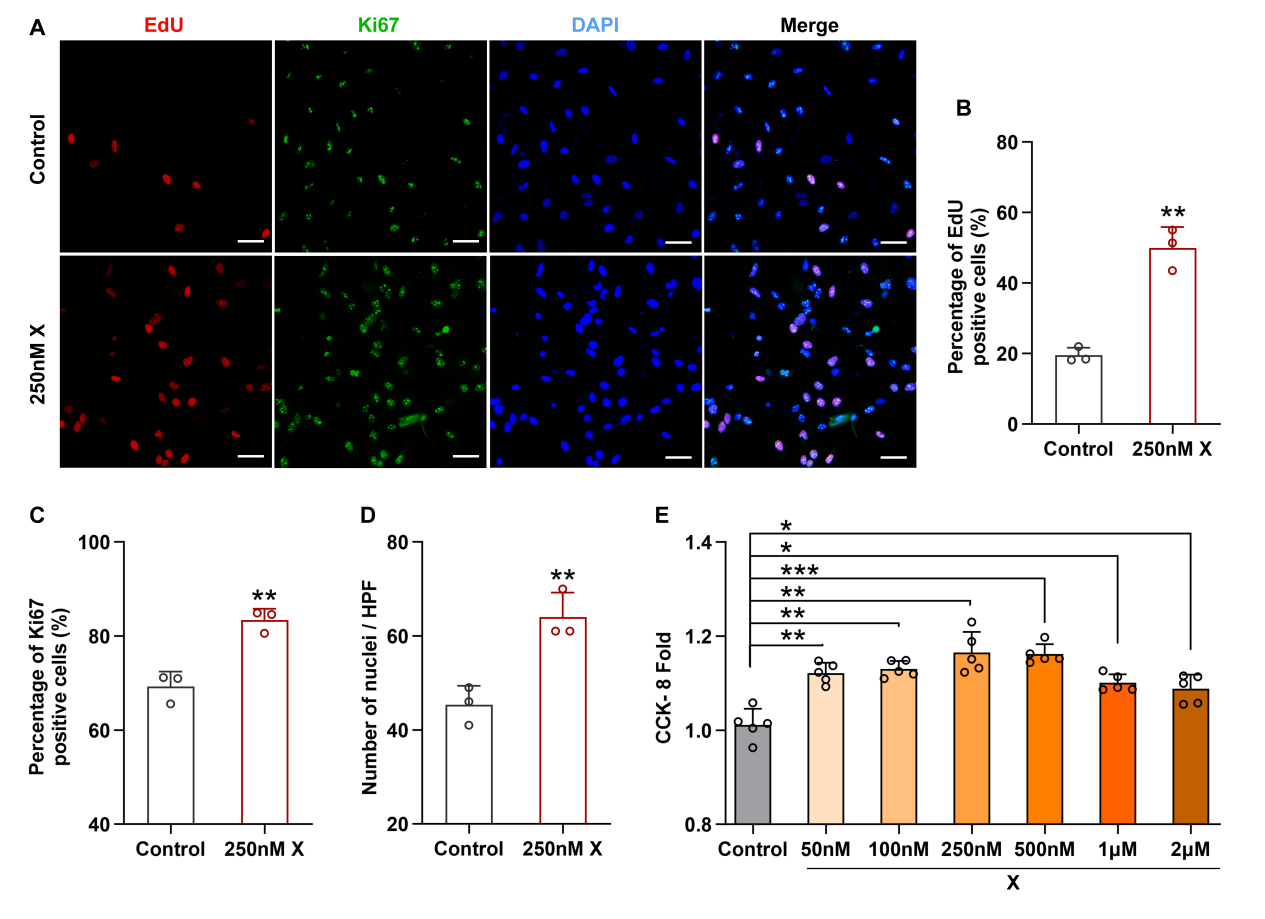

**Figure S5. XMU-MP-1 promotes proliferation of B4G12 cells.** (A) Representative EdU staining and immunofluorescence of Ki67 in B4G12 cells. (B to D) Percentages of EdU (B) and Ki67 (C) positive cells and nuclei counting (D) in (A) (n=3). (E) CCK-8 assay of B4G12 cells (Fold of control, n=5). Three visual fields were randomly selected in each sample for statistical processing. *P < 0.05, **P < 0.01, ***P < 0.001, error bars, mean ± SEM. Scale bars represent 50 µm. X, XMU-MP-1.

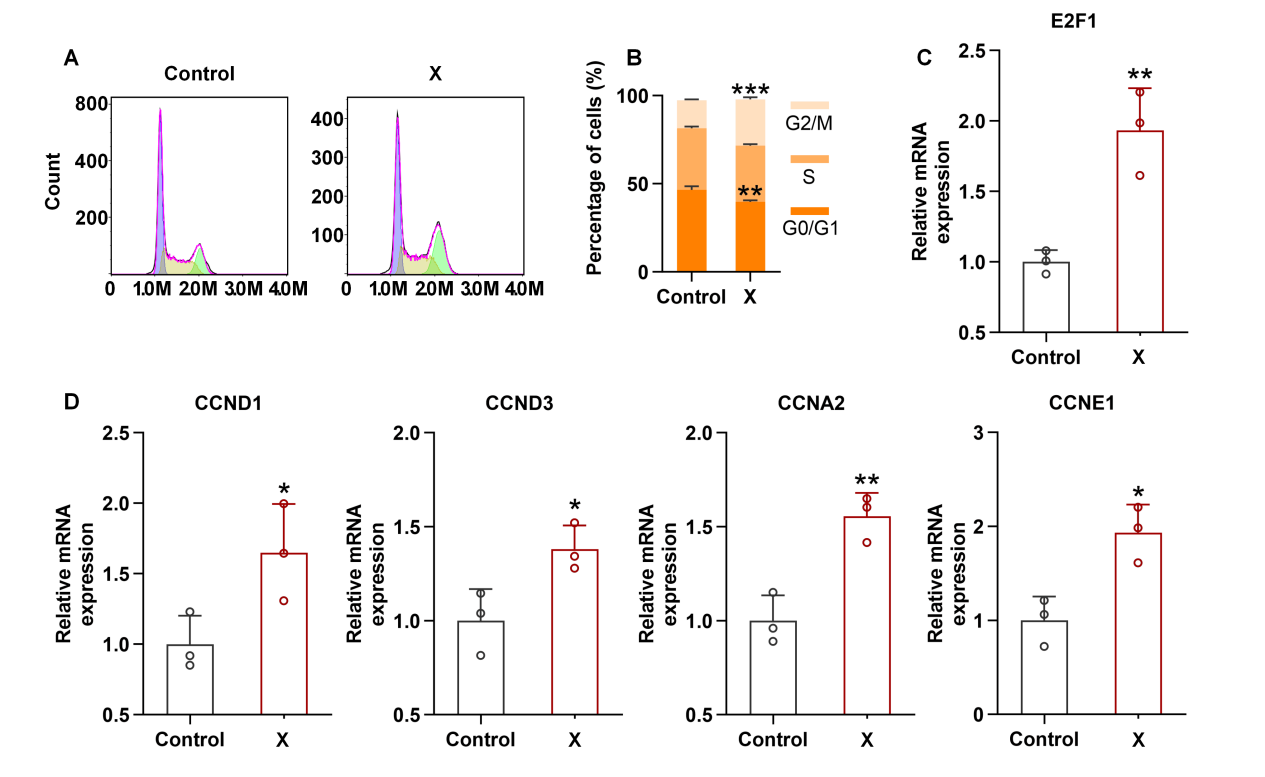

**Figure S6. XMU-MP-1 promotes cell cycle processes in primary rabbit CECs.** (A) Counts of G1, S, and G2/M-phase rabbit CECs treated with XMU-MP-1. (B) Percentages of G1, S, and G2/M phase rabbit CECs. (C, D) Relative mRNA levels of E2F1, CCND1, CCND3, CCNA2, CCNE1 in primary rabbit CECs. *P < 0.05, **P < 0.01, ***P < 0.001, error bars, mean ± SEM. X, XMU-MP-1.

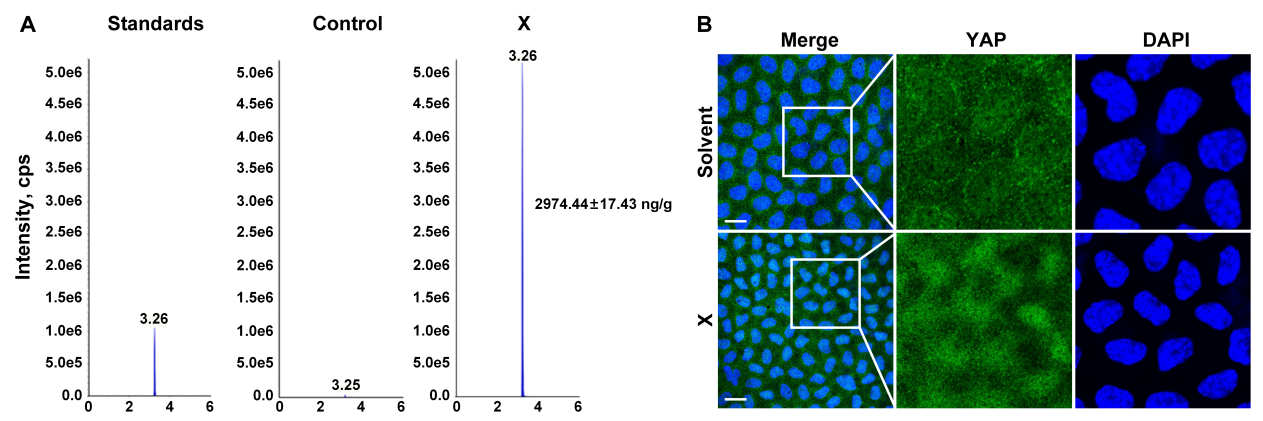

**Figure S7. Corneal endothelium absorbed XMU-MP-1 after ocular surface topical application.** (A) The concentration of XMU-MP-1 detected by LC-MS in the corneal endothelium of rabbits after topical application of XMU-MP-1 (50μM) for 3 days, with solvent as control. (B) Representative immunofluorescence images of YAP in mice corneal endothelial whole mount after topical application of XMU-MP-1 (1μM) for 3 days. Scale bars represent 10 µm. X, XMU-MP-1.

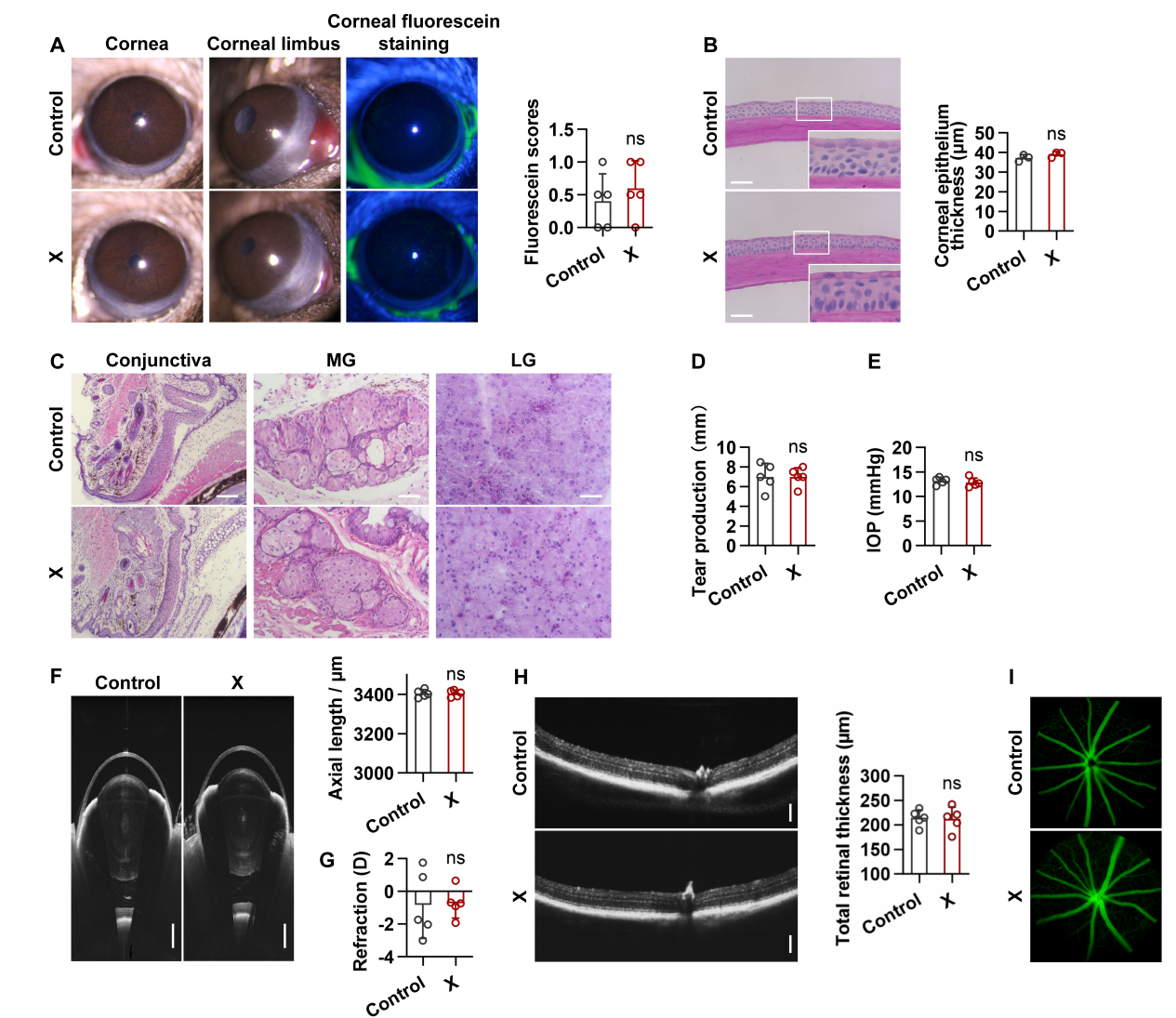

**Figure S8. XMU-MP-1 exhibits ocular safety profile in mice.** (A) Representative slit-lamp images of mice corneas, corneal limbus andcorneal fluorescein staining, as well as the fluorescein scores (n = 5). (B) The representative H&E staining images of mice corneas and the corneal epithelium thickness (n = 3). (C) Representative H&E staining images of mice conjunctiva, Meibomian gland and lacrimal gland. (D) Tear production of mice (n = 5). (E) IOP of mice after topical administration of XMU-MP-1. (F) Whole-segment OCT images and the axial length of mice eyes. (G) Refraction of mice eyes. (H) Retinal OCT images and total retina thickness of mice. (I) Representative fundus fluorescein angiography images of mice after XMU-MP-1 treatment. ns, no significance, error bars, mean ± SEM. Scale bars represent 500 µm in (H), 200 µm in (F), and 20 µm in (B and C). X, XMU-MP-1.

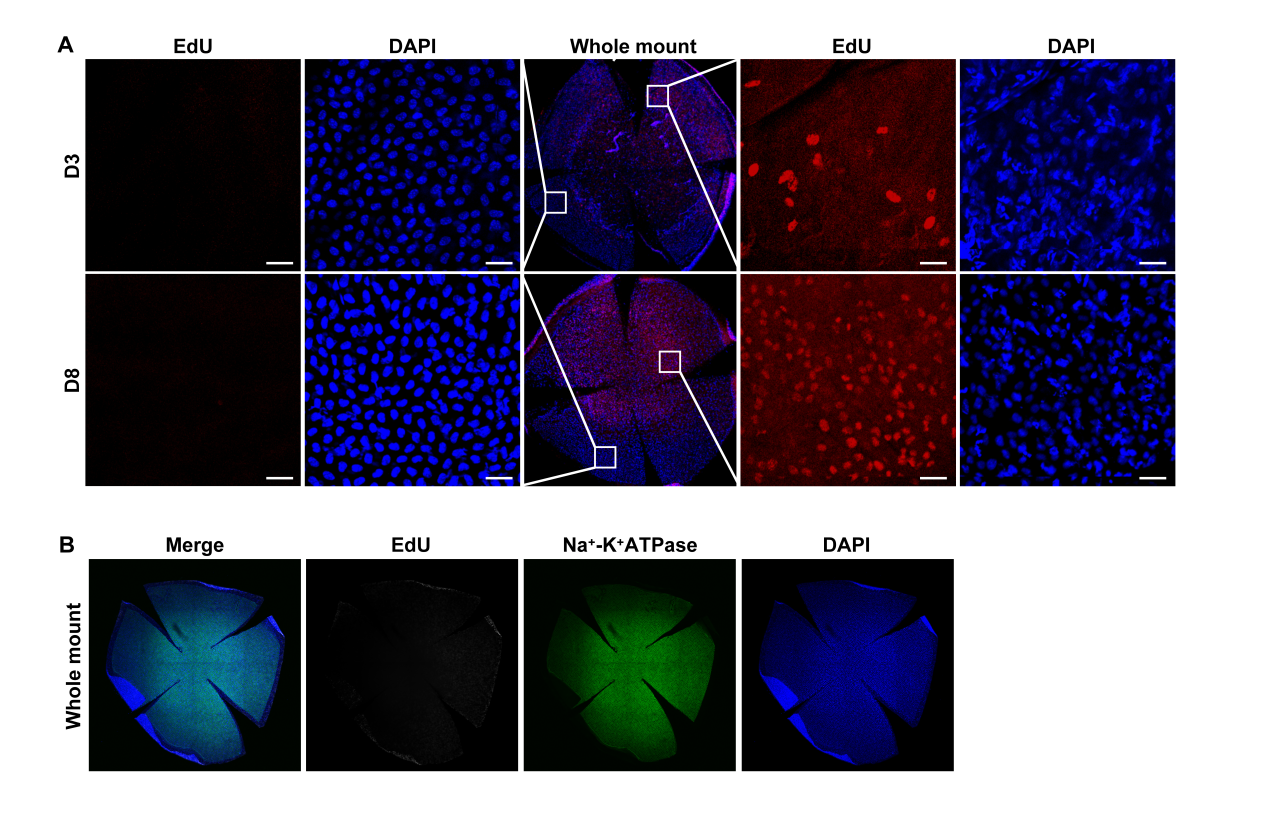

**Figure S9. Localization of EdU-positive cells during mouse corneal endothelial wound healing.** (A) Representative EdU staining of mice corneal endothelium during wound healing. (B) Representative EdU staining and immunofluorescence of Na^+^-K^+^ ATPase in normal mice corneal endothelium after intraperitoneal injection of EdU (50 mg/kg) for 7 days. Scale bars represent 10 µm.

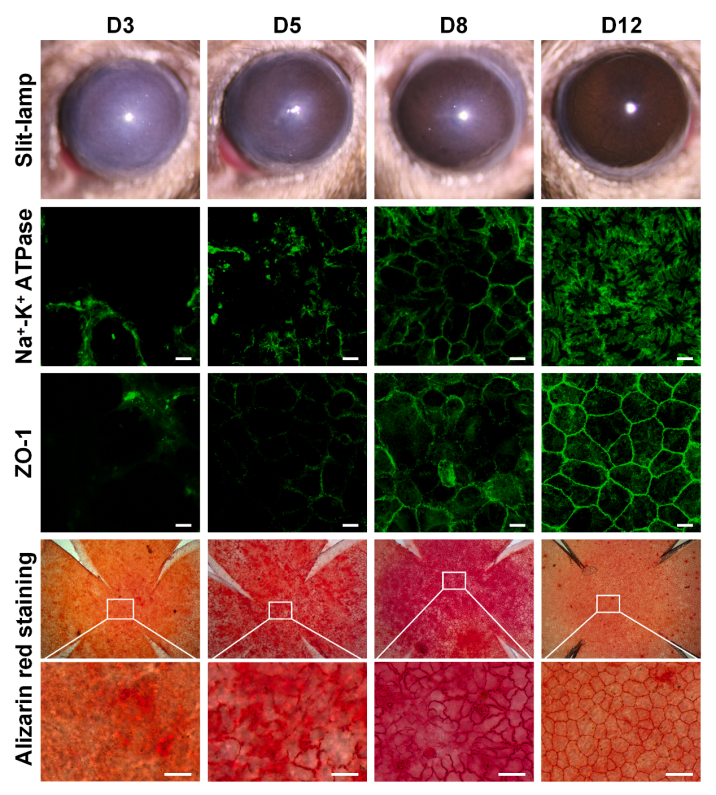

**Figure S10. Changes in cell morphology and function during mouse corneal endothelial wound healing.** Representative slit-lamp and alizarin red staining images and immunofluorescence of ZO-1 and Na^+^-K^+^ ATPase in mice corneal endothelial whole mount during wound healing. Scale bars represent 10 µm.
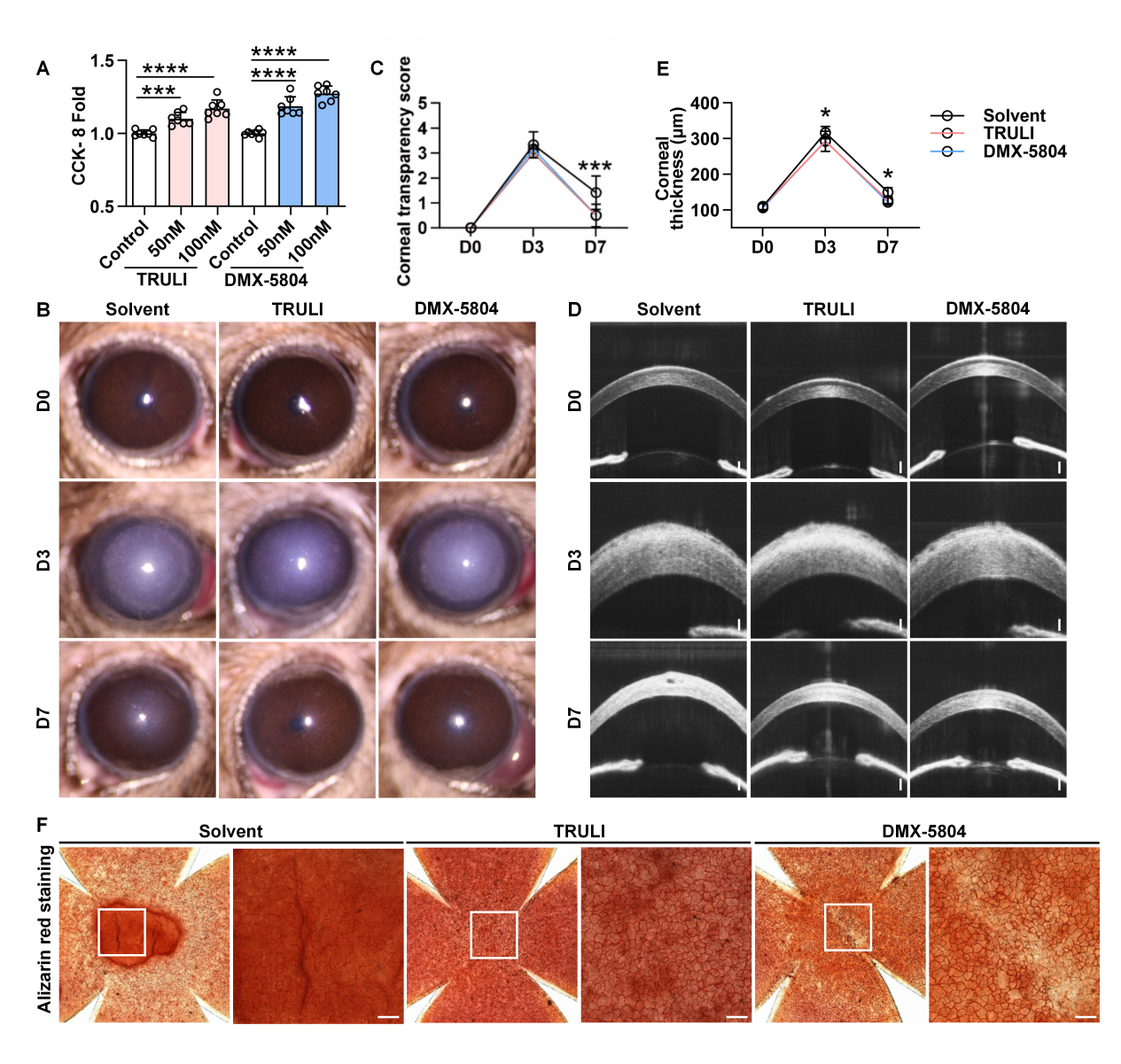

**Figure S11. TRULI and DMX-5804 promote corneal endothelial wound healing in mouse model.** (A) CCK-8 assay of primary rabbit CECs treated with DMSO, TRULI and DMX-5804 (Fold of control, n = 7). (B to E) Representative slit-lamp images (B) and anterior segment OCT images (D) of mouse model treated with TRULI and DMX-5804. Corneal transparency score (C) and central corneal thickness (E) of mice shows TRULI and DMX-5804 promoted recovery of corneal opacity and edema (n = 6). Corneal alizarin red staining (F) demonstrated that TRULI and DMX-5804 promotes wound healing of CECs in mice. *P < 0.05, **P < 0.01, ***P < 0.001, error bars, mean ± SEM. Scale bars represent 100 µm in (D) and 40 µm in (F).

**Table S1. AAV sequences**

| **Target** | **Forward** | **Reverse** |
| --- | --- | --- |
| AAV Yap1 shRNA | ACTTGGAGGCGCTCTTCAATGCCGTCATG | CATGACGGCATTGAAGAGCGCCTCCAAGT |

**Table S2. siRNA sequences**

| **Target** | **Forward** | **Reverse** |
| --- | --- | --- |
| Rabbit YAP siRNA | GAGAUACUUCUUAAAUCAC | GUGAUUUAAGAAGUAUCUC |
| Rabbit NC siRNA | UUCUCCGAACGUGUCACGUTT | ACGUGACACGUUCGGAGAATT |
| Human YAP siRNA | GACCAAUAGCUCAGAUCCUUU | AAAGGAUCUGAGCUAUUGGUC |
| Human NC siRNA | UUCUCCGAACGUGUCACGUTT | ACGUGACACGUUCGGAGAATT |

**Table S3. qPCR primers**

| **Target** | **Forward 5’-3’** | **Reverse 5’-3’** |
| --- | --- | --- |
| *Ccne1* | GGTGTCAGGGTATCAGTGGTGTG | TGGCTTTCTTTGCTTGGGCTTTG |
| *Ccnd3* | TGCGGAAGATGCTGGCGTAC | GACAGGTAGCGATCCAGGTAGTTC |
| *Ccnd1* | CAAGCAGATCATCCGCAAGC | GGTAGCAGGACAGGAAGCTG |
| *E2f1* | AGGACCTTCGGAGCATTGTG | ACATCGATGGGGCCTTGTTT |
| *Ccna2* | CCCTTGTCTCGTGGACCTTC | GTGTCTCTGGTGGGTTGAGG |
| *Yap1* | CCAAGGCTGGACCCTCGTTTTG | TTGCTGCTGCTGGTTGGAACTG |
